# A developmental stretch-and-fill process that optimises dendritic wiring

**DOI:** 10.1101/2020.07.07.191064

**Authors:** Ryan Rahy, Lothar Baltruschat, André Ferreira Castro, Peter Jedlicka, Gaia Tavosanis, Hermann Cuntz

## Abstract

Circuit connectivity and computation depend on how dendrites branch and occupy space within neural tissue. While optimal wiring principles have long been known to constrain dendritic morphology and their scaling behaviour, the growth dynamics that produce such optimised structures remain unclear. Leveraging structural imaging across development, we identify two complementary growth strategies – inside-out versus outside-in – that together generate mature dendritic arbours. We formalise these dynamics in a mathematical model that captures the two growth modes and show that their interplay yields wiring-efficient, space-filling morphologies and class-specific developmental trajectories across species. This framework provides an algorithmic account of how local branching dynamics give rise to globally optimised architectures. By linking dendritic growth rules to functional design constraints, our theory offers a unifying description of dendritic differentiation and a basis for understanding how coverage and connectivity emerge during neural circuit formation.

**In brief:** We derive a detailed mathematical model that describes long-term time-lapse data of growing dendrites; it optimises total wiring and space-filling.

**Highlights:**

- Fly neurons stretch and fill a given target area with precise scaling relations.
- We observe a sequence of two growth strategies.
- Each growth type implements optimal wiring which leads to optimal space filling.
- A model combining these programs captures the development of dendritic structures.

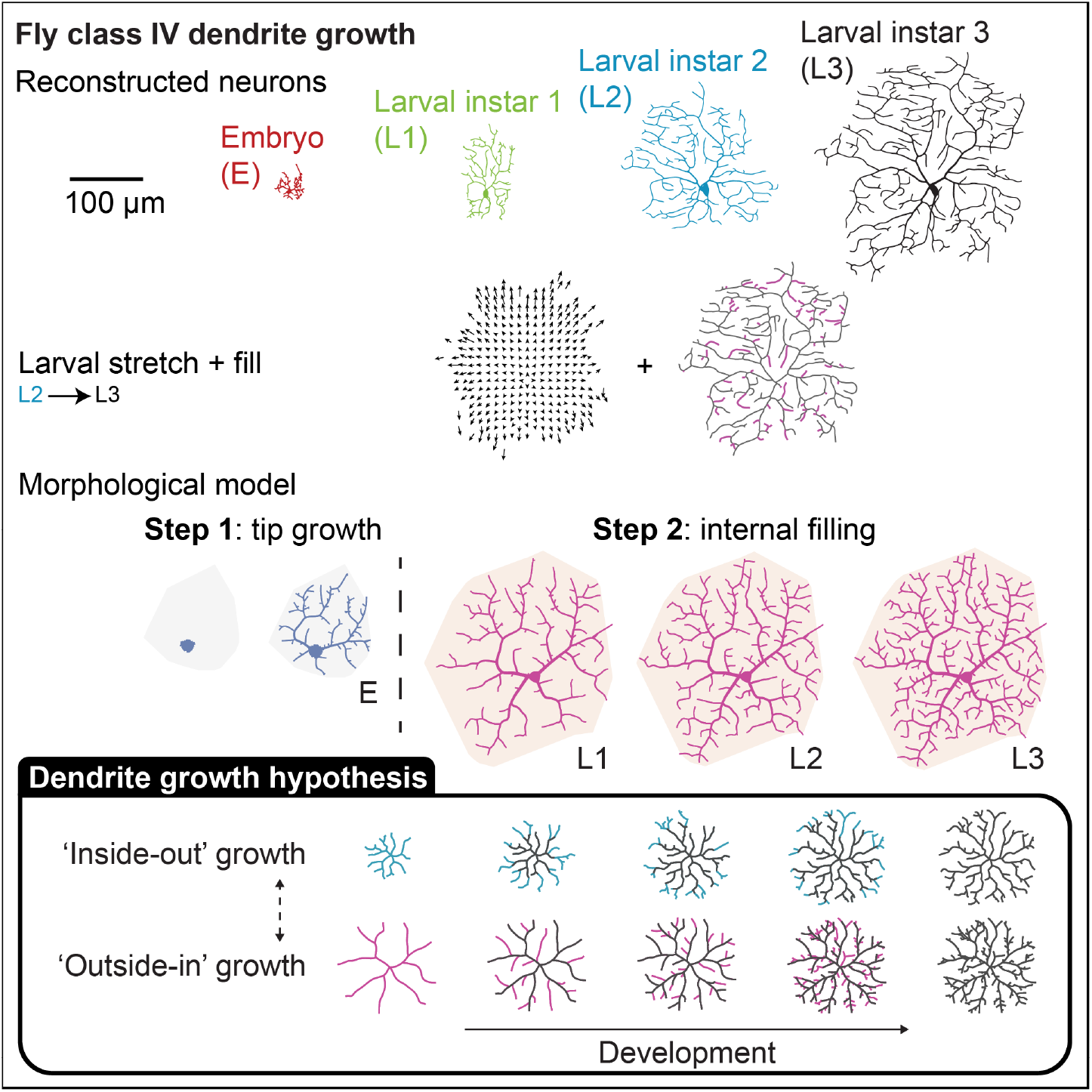

## Introduction

Dendritic branching defines the structural and functional identity of neurons by shaping how they receive, integrate, and transmit information within neural circuits (Jan and Jan, 2003; London and Häusser, 2005; Poirazi and Papoutsi, 2020). Through complex arborisation patterns, dendrites provide the receptive surface that underlies the brain’s high degree of connectivity, ultimately defining the connectome (Winding et al., 2023; Dorkenwald et al., 2024; Schneider-Mizell et al., 2025; Piazza et al., 2025; Fritz et al., 2025). Class-type specific morphologies emerge from coordinated influences of genetic programs and growth-promoting signals that activate local interactions guiding branches into their appropriate spatial domains (Parrish et al., 2007; Lefebvre et al., 2015; Barabási et al., 2025). Although these molecular and cellular regulatory processes have been extensively characterised, the branching dynamics that govern dendritic organisation remain poorly understood.

While several computational approaches can generate dendrite-like branching structures (Ascoli et al., 2001; Luczak, 2006; Koene et al., 2009; Cuntz, 2015; Kanari et al., 2022; Kirchner et al., 2025), morphological models based on optimal wiring (Cuntz et al., 2007, 2008; Budd et al., 2010; Cuntz et al., 2010) are particularly well suited for studying dendrites and axons in their role to implement circuit connectivity. Models derived from minimum spanning trees (MSTs) (Kruskal, 1956; Prim, 1957) successfully reproduce key geometric and topological features of dendritic architecture and provide a normative explanation for conserved scaling relationships among neuronal size, wiring length, and connectivity (Cuntz et al., 2012). However, as MST-based models describe what optimal wiring looks like, they leave open the question of how branching dynamics in neurons achieve this during development.

Recent advances in time-lapse imaging now make it possible to observe dendritic growth as a continuous process rather than a sequence of static morphologies (Sin et al., 2002; Lee et al., 2013; Hogg et al., 2021). Quantitative live imaging captures branch extension, retraction, and remodelling with sufficient temporal resolution to test computational models of arbour formation (Mizrahi, 2007; Lee et al., 2013; Chalmers et al., 2016; Gonçalves et al., 2016; Radic et al., 2017; Ferreira Castro et al., 2020; Stürner et al., 2022; Palavalli et al., 2021; Shree et al., 2022; Rigaux et al., 2025). These studies reveal that class-specific dendritic arbours emerge through sequential growth stages combining stochastic branch exploration with deterministic refinement (Ferreira Castro et al., 2020; Palavalli et al., 2021). Despite these advances, imaging alone cannot explain how local growth dynamics give rise to globally optimised architectures.

To uncover the growth rules that generate optimally wired dendritic morphologies, we focused on class IV dendritic arborisation (da) neurons of *Drosophila* larvae, a well-established model for dendrite morphogenesis (Jan and Jan, 2010; Grueber et al., 2002). These sensory neurons form a highly branched, space-filling meshwork that tiles the larval body wall through local self-avoidance and interactions with neighbouring arbours (Sugimura et al., 2007; Matthews et al., 2007; Soba et al., 2007; Shimono et al., 2010; Shree et al., 2022). Their planar geometry, stereotyped architecture, and genetic accessibility enable direct quantification of branch dynamics during growth, making them an ideal system for extracting general rules of dendritic self-organisation.

To formalise dendritic maturation, we develop here a developmental morphological model grounded in experimentally-observed growth dynamics after tracking all branches of class IV da neurons (c4da) during development. The model reconstructs developmental trajectories from quantitative morphometrics, incorporating two complementary growth programs: an initial inside-out expansion driven by tip extension and a secondary outside-in phase that fills space through new branch formation and stretching. This dual-process framework reproduces class-specific morphologies and explains how neurons achieve optimal wiring while occupying available territory. Validation of our model based on morphological statistics using synaptic-resolution, densely reconstructed datasets confirms that the local growth dynamics that we describe generalise across two- and three-dimensional environments and across species. Together, these results establish a unified generative model of dendritic development that links local growth rules to global wiring optimisation.

## Results

### Scaling behaviour of C4da neurons throughout larval development

In order to dissect how optimal wiring is implemented in the growth of individual cells, we used live imaging to obtain time-lapse reconstructions of developing dendrites of the same C4da neurons throughout larval development (embryonic stage E – *red*, and instar larval stages L1 – *green*, L2 – *blue*, L3 – *black*, **Figure 1A**, see **Methods** “Time-lapse image acquisition”). Remarkably, large parts of the dendritic architecture were conserved over all stages of development while the overall shape was stretched, increasing the size in all directions (**Figures 1B and C**). Most importantly, individual dendritic branches from previous stages were easily identifiable and new branches grew in the space that became available through the extensive epidermal stretching.

**Figure 1.**
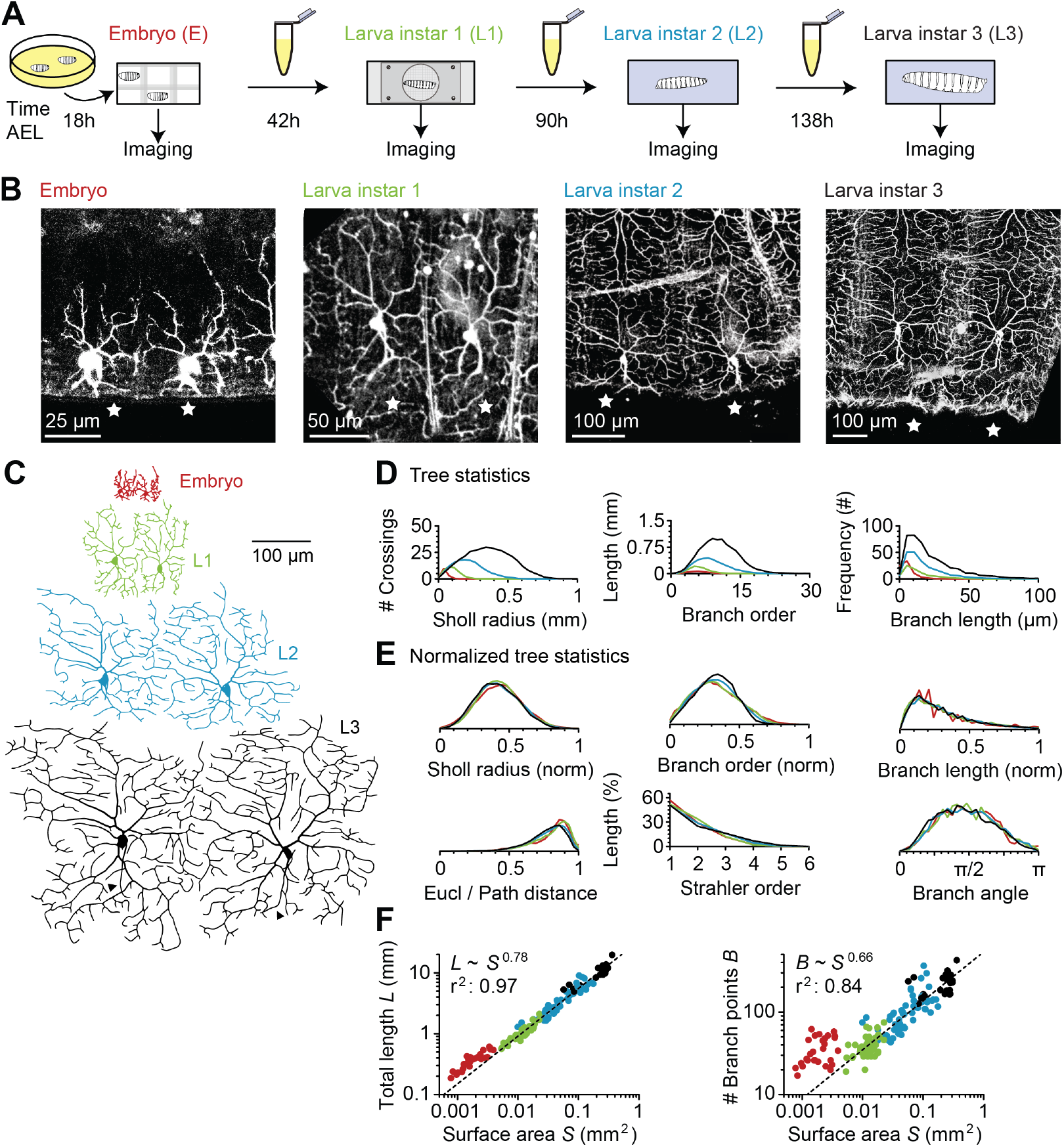
Long-term time-lapse imaging of C4da neuron development. **A**, Diagrams illustrating the imaging procedure throughout *Drosophila* larva developmental stages: embryo (E), larval instar (L) 1, 2 and 3. The larvae were kept in culture medium between the imaging steps. Times shown are AEL (after egg laying). **B**, Maximum intensity projections of image stacks from the same two neighbouring da dendrites (stars) for all four developmental stages. **C**, Digital reconstructions of the two neighbouring dendrites from **B** with small diameters clamped at 2*µm* for better visualisation. Triangular markers in L3 show axons, which are ignored in the rest of the study. **D–E**, Average branching statistics of reconstructions at the four developmental stages in absolute (**D**) and normalised (**E**) values. In all panels, the colours for the developmental stages are E (red), L1 (green), L2 (blue), L3 (black). **F**, Scaling behaviour of total length *L* and number of branch points *B* against the total surface *S* that the dendritic trees span. Dashed lines show the best power law fits (linear fit in the log-log domain) for the data points from L1–L3 with respective power and *r*^2^ values.

We looked at different dendritic morphological parameters throughout the E–L3 larval stages to better quantify the geometrical aspects of the growth process. All branching statistics scaled up extensively with the developmental stages (**Figure 1D**) but were essentially invariant of the developmental stage once the differences in overall sizes were normalised out (**Figure 1E**), suggesting a general growth process conserving dendritic features.

The surface area *S* covered by the dendritic trees increased by a factor of 62× between embryo (E) and third instar larva (L3) stages. The relation between the surface area *S* and the total dendrite cable length *L* during the L1–L3 stages was consistent with a strict power relation of exponent 0.78 (*L* ∼ *S*^0.78^; *R*^2^ = 0.97). This implied that branches filled up newly available space more than in a simple stretching process (*L* ∼ *S*^1*/*2^, increasing the size without adding branches) but without conserving cable density (*L* ∼ *S*, a linear relation with an exponent of 1) throughout development (**Figure 1F**, *left panel*). Similarly, the number of branch points *B* scaled with the surface area with an exponent of 0.66 (*B* ∼ *S*^0.66^; *R*^2^ = 0.84), reflecting a branching process falling somewhere between a pure stretching process with no new branches (*B* ∼ *S*^0^, i.e. *B* does not change with *S*) and a process conserving branch point density (*B* ∼ *S*; **Figure 1F**, *right panel*). The embryonic stage, on the other hand, showed markedly different scaling relations between these morphological parameters compared to L1–L3, suggesting that C4da neurons might exhibit a different initial growth process (**Figure 1F**), in line with previous observations (Parrish et al., 2009). Overall, the analyses suggest a simple progression of growth with strong quantitative constraints for our mathematical theory.

### Dendritic developmental stretch-and-fill

Beyond the quantification of the mathematical scaling relations, we dissected the C4da developmental growth process, leveraging our detailed time-lapse data to compare each time point to the next. We tracked branch and termination points that were conserved in consecutive reconstructions using a manual registration procedure (**Figure 2A**, see **Methods** “Time-lapse analysis”). The analysis unveiled a surprisingly isometric stretching (**Figure 2B**) with a tight linear slope between the stretching vector and the distance of points from the dendrite root (**Figure 2C**). Interestingly, the overall origin of stretching did not lay precisely in the dendritic root location (**Figure 2B**), consistent with a possible impact of superficial epithelium extension on dendritic stretching during development (Jiang et al., 2014). The results from our manual registration were all confirmed by an automated tracking of branches (see supplementary **Figure ??**, Hermila et al., 2026, in preparation).

**Figure 2.**
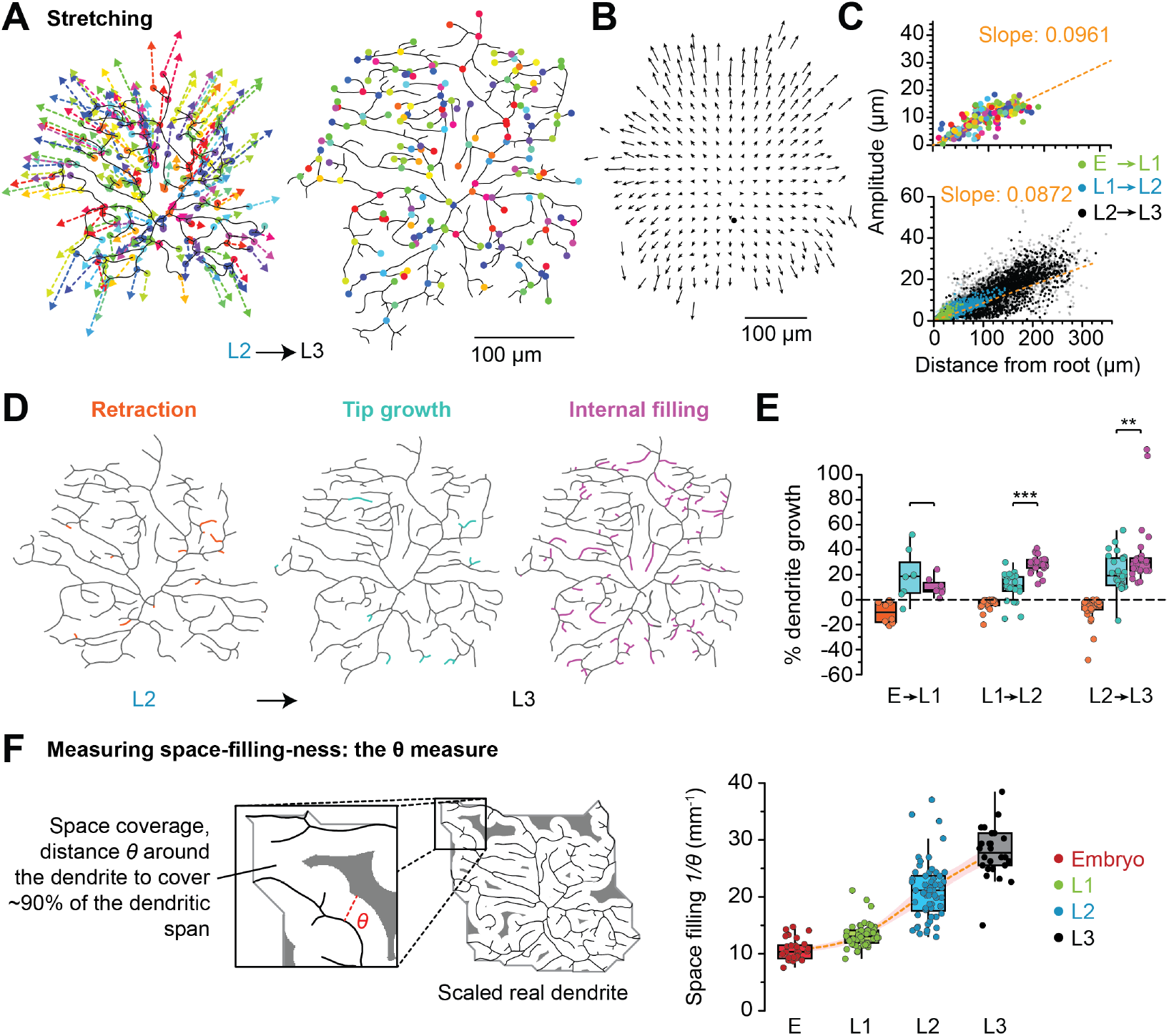
C4da neuron developmental stretching. **A**, Time-lapse data analysis: Branch and termination points (coloured dots) in time 1 (*left*, here L2) are registered to the corresponding points in time 2 (*right*, here L3, same colours). The arrows on the left depict the movement of the respective topological points and illustrate the stretching of the entire structure between two acquisitions. **B**, Overall analysis for all pairs (*N* = 47). Arrows (every 25*µm*) are binned averages only shown where data for at least 5 pairs was available. Dendrites were centred on their root (black dot). **C**, Analysis of stretching by comparing the arrow lengths (*top*, one dot per topological point with same colours as in **A**; *bottom*, topological points collected from all pairs with colours indicating larval instar at time 2 and their distance to the root). Greyed out points represent tips, linear fits (dashed orange lines) determine the slopes. **D–E**, Retraction (orange), tip growth (cyan), and internal filling (purple) processes observed in C4da cells, illustrated in **D** and quantified across all structures and phases in **E. F**, *Left*, diagram showing how *θ* is calculated as the distance one has to move away from the tree to cover ∼ 90% of the spanning field of the dendrite, an inverse of space-filling-ness for neurons. *Right*, calculation of *θ* for all stages. Dashed orange line shows a smoothed Eilers estimate of the data. In panels **D–F**, all dendritic morphologies were scaled to cover a reference surface area of (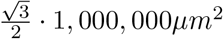, see **Figure ??**).

Having quantified the overall stretching of the dendritic spanning field, we scaled all the dendritic reconstructions to the same surface area to focus on the changes inside. This was done after applying a combination of small global alignment steps (a rotation with a consecutive stretching in both spatial dimensions) to superimpose the obvious conserved branching patterns. Our time-lapse registrations enabled us to then characterise the precise changes in the branches that were not conserved. We quantified the branch dynamics and separated them into: (1) retraction (branch elimination), (2) tip growth (branch elongation/shrinkage), and (3) internal filling (new branch formation) (**Figure 2D**). Quantifying each of these types of changes revealed that the amount of branch retraction was low and decreased throughout development. In addition to that, while E to L1 showed more tip growth than internal filling, this trend reversed for L1 to L3 that showed more internal filling than tip growth, once again indicating a difference in developmental program between the embryonic and larval stages (**Figure 2E**).

The considerable amount of internal filling and tip growth inside the dendritic spanning area may be linked to the space-filling property of C4da neurons. We therefore introduce a measure for the efficient coverage of available surface area *S*: *θ*, the maximum distance one needs to move away from the dendrite to cover ≈ 90.69% of the dendritic span. This measure of space filling is inspired by considerations of circle packing on a hexagonal grid (**Figure 2F**, see **Methods** “Space-filling measure 1*/θ*” and supplementary **Figure ??**). Its inverse, 1*/θ*, then quantifies how well the neuron fills the dendritic span (**Figure 2F**, *right*). In the current case where all morphologies were stretched to cover the same surface area *S*, 1*/θ* increased steadily during larval instar growth L1–L3.

### Morphological model of dendrite development

Having quantified the simple progression of growth iterations and scaling of branching statistics in the developmental dataset, we turn to morphological models of neurons to analyse the growth through the lens of optimal wiring. We applied a minimum spanning tree (MST) algorithm that optimises total cable length on our current C4da reconstructions data (Cuntz et al., 2007; Prim, 1957, see Supplemental section “MST-based model reproducing C4da development”). This showed that C4da dendrites followed optimal wiring principles throughout development and were best matched with a trade-off parameter *bf* = 0.225 between minimising total dendrite length *L* and path lengths *PL* along the tree from target points toward the root (*wiring cost* = *L* + *bf* · *PL*; **Figure ??**; see also Cuntz et al., 2010; Wen and Chklovskii, 2008; Nanda et al., 2018). We next explored how the developmental process leads to such optimally-wired trees from one stage to the next.

To this end, and based on our analysis of C4da development, we put forward a general hypothesis of dendritic development, with two main processes in which dendrites spread into the available space around them (**Figure 3**). Inside-out growth describes dendritic changes that occur at the tips of the dendrites, with more tip retraction and elongation and minimal interstitial branching; Outside-in growth describes a spreading pattern that fills out the available spanning field with longer, relatively sparser primary dendrites before filling the gaps in with interstitial branches. The resulting dendritic spreading pattern of a given structure can involve both growth processes to varying degrees.

**Figure 3.**
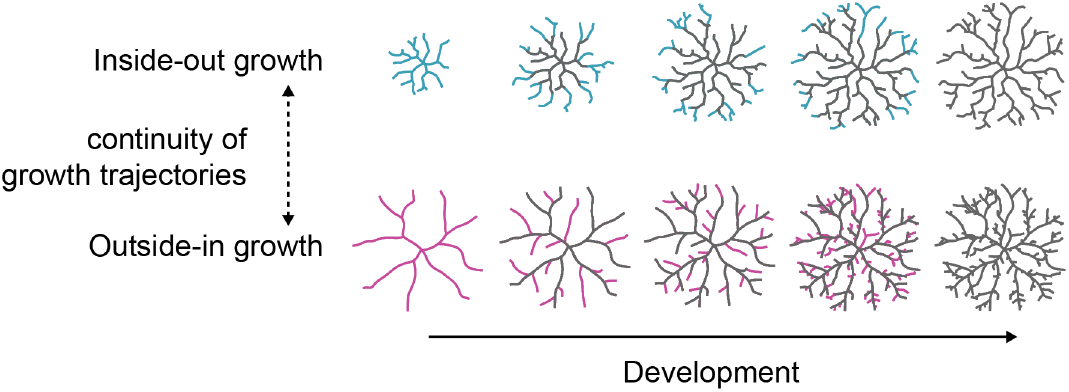
Dendritic growth hypothesis. Model incorporating inside-out growth (*top*) and outside-in growth (*bottom*), with new branches at every developmental step in *light blue* and *magenta* respectively compared to the conserved tree from previous developmental time points (in *grey*).

We formalised this hypothesis in an algorithm that builds on optimal wiring principles and can generate different developmental trajectories. Developmental dynamics are determined by a new parameter, the spreading pattern *sp*, which controls the timing of emergence of different outgrowths (**Figure 4A**, *bottom panels*). The resulting temporal patterns can fall anywhere on the continuous spectrum between the two growth types, with lower *sp* values favouring inside-out growth and higher *sp* values favouring outside-in growth. Similar to the aforementioned MST-based model, the branching pattern of the modelled structures is determined by a balancing factor *bf* (**Figure 4A**, *top panel*). The resulting branching structure is independent of the *sp* value, as can be seen in the distinct centripetal biases of the root angle distributions (calculated as the distribution of angles between each dendritic segment and its direct path to the soma Bird and Cuntz, 2019, **Figure 4B**, *bottom panel insets*) of the model structures; this suggests consistent wiring optimality across *sp* values. In addition to that, the scaling of the total dendritic length *L* with space filling 1*/θ* was conserved throughout the parameter space of our model (**Figure 4C**).

**Figure 4.**
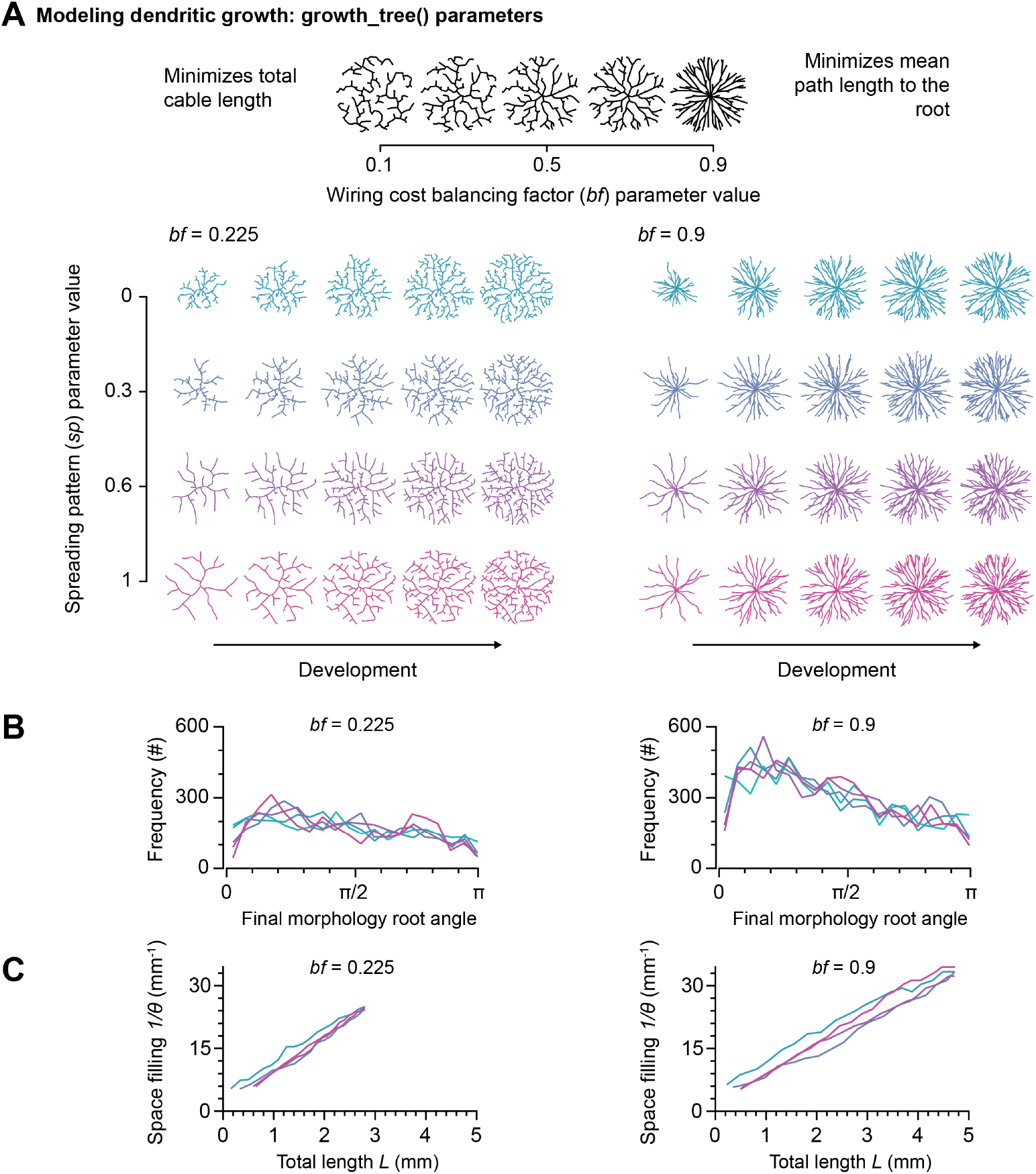
growth_tree() algorithm is based on optimal wiring principles enabling distinct developmental trajectories along the continuous spectrum between the inside-out and outside-in growth type. **A**, Effect of the main parameters on the output of the proposed algorithm modelling dendritic growth. *Top row*, Effect of changes in balancing factor (*bf*) for a set *sp* = 0. *Bottom panels*, Effects of changing the spreading pattern parameter *sp* for *bf* = 0.225 (*left*) and *bf* = 0.9 (*right*) on the developmental iterations of growth. **B**, Root angle distributions of the final morphologies from the bottom panels of **A. C**, Total length *L* vs space filling 1*/θ* (see **Figure 2F**) during the developmental trajectories shown in the bottom panels of **A**.

In order to generate realistic structures, an additional parameter *k* injects noise into the growth process, subtly affecting the resulting structures along with their wiring cost optimality and growth trajectories (**Figure ??**). **Figure ??** compares the growth dynamics and scaling behaviours of these optimal-wiring-based morphological models.

### Morphological model of C4da neurons throughout development

With this growth model, we simulated the complete *de novo* development of C4da dendrites in two steps in order to capture the change in growth program described earlier (**Figure 5A**). We first modelled the embryonic structure with a low *sp* = 0.3 then continued development until L3 with a higher *sp* = 1 to simulate the change from the inside-out growth program to outside-in. The *bf* parameter was unchanged from the wiring-optimal MST models (**Figure ??A**, *left*), and remained consistent throughout development. The synthesised L1 and L2 structures were selected as the intermediate model structures that length-matched the reconstructed structures.

**Figure 5.**
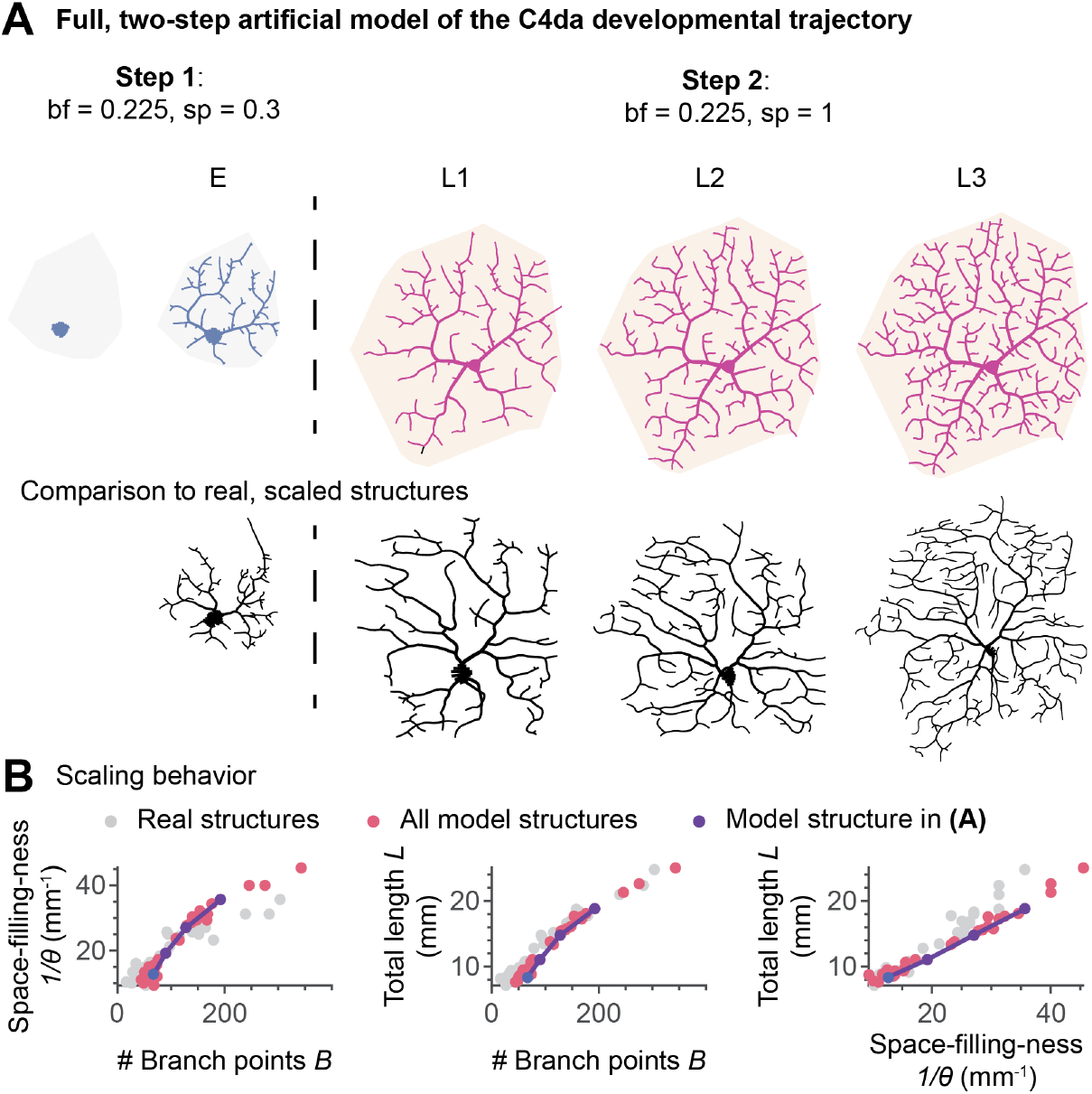
Model of C4da development. **A**, A two-stage model of the developmental trajectory of C4da cells (*top*), compared to the reconstructed structures scaled appropriately (*below*). In both modelling steps *bf* = 0.225, while *sp* was changed from 0.3 to 1 between the E and L1 stages. The shaded area behind each neuron shows the real dendritic span used to grow these neurons. Synthesised structures are coloured according to the *sp* value, following the colour scheme in **Figure 4A. B**, Scatter plots showing the scaling behaviour of real dendrites (in grey) compared to model dendrites (in pink), with the model trajectory from **A** shown in purple. Note that only structures for which all developmental phases were available were used.

Applying this model to all real C4da structures reproduced the scaling behaviour that real dendrites underwent while optimising wiring and space filling throughout development (**Figure 5B**). Modelling C4da cells in a single step without the switch in growth programs led to structures with imbalanced branch lengths (see Supplemental section **??**).

### Applying the growth model to other cell types

Our developmental growth process model reproduced the morphologies and the space-filling characteristics of other example dendrites, both planar (fly C3da sensory neurons and lobula plate tangential cells) and three-dimensional (rat dentate gyrus granule cells, fly second order nociceptive neurons, and rat cortical layer 5 pyramidal cells), with *bf* and *sp* parameter combinations that spanned different parts of the parameter space (**Figure 6**). The structures predicted for these neurons were confirmed by statistical comparison to reconstructed neurons (**Figure ??**).

**Figure 6.**
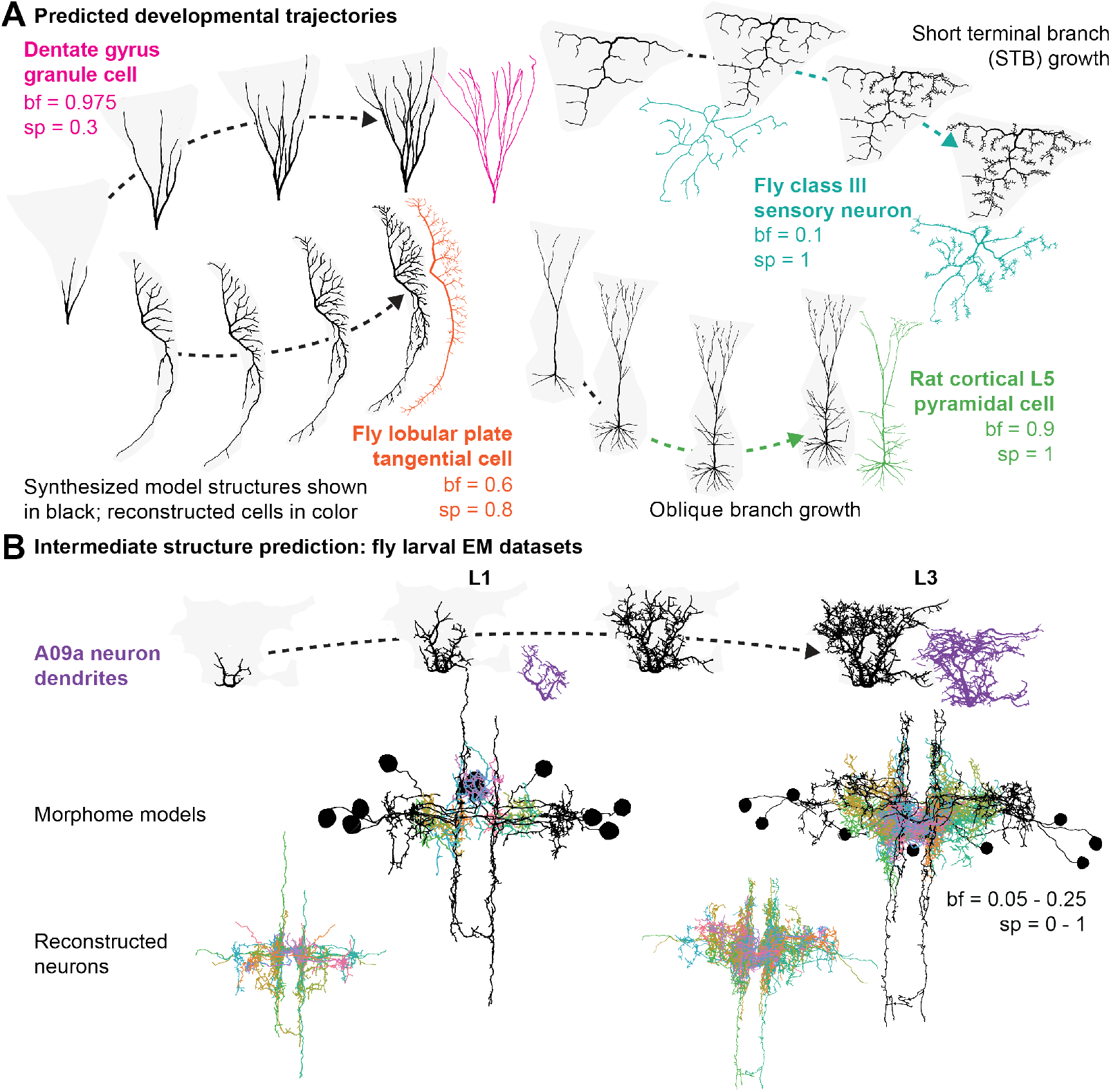
Modelling the development of additional cell types. **A**, Growth model applied to a variety of neurons: *Orange*, Fly tangential cell; *pink*, Rat granule cell; *green*, Rat cortical pyramidal cell; *teal*, Fly C3da neuron. Synthesised structures from the model are shown in black and real cells in colour, with the grey shaded area marking the spanning area (derived from the reconstructed trees) in which these dendrites were grown. **B**, Intermediate structure prediction paradigm: *Top*, an example growth trajectory of a fly Basin A09a neuron (*lila*) reconstructed from EM data at larva instar stages L1 and L3. Synthesised L3 structure and its intermediate growth structures (*black*). *Bottom*, connectomes of all nociceptive neurons with the real axons and linking cables (in *black*) and modelled (large images) and real (insets) dendritic structures. Each colour represents a different dendrite.

Similar to C4da cells in **Figure 5A**, modelling the entire development of some other neuron types required multiple steps (**Figurs 6A**). This included dendrites with growths having differential spatial organisation, such as short terminal branch growth in C3da cells, and growths with differential timing, such as oblique branch growth in L5 cortical pyramidal neurons.

In order to further validate the model developmental trajectories, we compared the intermediate structures generated by our model to the earlier stages of the biological counterparts. This was done using L1- and L3-stage structures of the same neurons obtained from two electron microscopy datasets of the same fly nociceptive circuit (Gerhard et al., 2017). In this paradigm, we first modelled the dendritic structures corresponding to L3 neurons, then analysed the intermediate model structures to evaluate how well our trajectories captured the properties of the reconstructed L1 structures. By length-matching the synthesised developmental steps to the reconstructed L1 structures (**Figures 6B**, *first row*), we show that our model was able to capture both the individual structures and the trajectories that give rise to them. To evaluate the overall performance across all models, we put together the artificial dendrite models at their original locations in the connectome data and re-inserted the original axons and soma-linking cables pertaining to the structures (**Figures 6B**, *second row*). Other than the expected difference in tissue packing and stretching at the L1 stage, the identified synthesised structures matched the real structures (**Figure S6**).

## Discussion

A central challenge in understanding neuronal development is to explain how complex dendritic arbours grow over time while maintaining functional organisation and wiring efficiency. The goal of this study was to obtain detailed time-lapse recordings of class IV dendritic arborisation (C4da) morphologies and comprehensively analyse developmental dendritic branching changes. We present a morphological model that captures the dendritic development constrained by that data as well as optimal wiring principles, and formulate a general hypothesis on dendritic growth.

From these analyses, several general principles of dendrite development emerge: (1) As the larva grows, C4da dendrites stretch with a remarkable conservation of branching structure and grow into new space as it becomes available; (2) Two alternative growth programs separated by what looks like a phase transition were observed throughout C4da development; (3) The two programs (inside-out vs outside-in) are required in the model to reproduce the data; (4) Both programs seem to guarantee optimal wiring throughout development; (5) Optimal wiring algorithms in general seem to guarantee that the dendritic span is optimally space-filled for a given dendritic cable length; (6) The growth model derived from C4da neurons applies widely to other neurons and explains existing data from cellular neuroanatomy.

From our *in vivo* time-lapse data we showed that the initial embryonic phase can be described as predominantly driven by external tip growth, consistent with previous observations (Shree et al., 2022). In contrast, the growth of the remaining developmental stages was better described by a regular and isometric stretching combined with internal in-filling (**Figure 2D,E**). This interpretation, of which we are confident due to the strong structural conservation between time points (see also our extensive appendix), is consistent with previous work showing a reorganisation during C4da development. One study found exocytotic machinery related to dendritic expansion shifting from being well-distributed throughout the entire dendrite during embryogenesis to becoming predominantly localised to primary (but not terminal) dendrites during larval development (Peng et al., 2015). Another study further showed that during development, C4da neurons first fill the empty space around them during embryonic development until they reach the hemisegment boundaries, then stretch loosely with the epithelium (Parrish et al., 2009). These changes in cellular mechanisms and extracellular contexts could reveal possible mechanistic or causal processes underlying the change in dendritic spreading patterns that we report here. Relatedly, the timing of similar changes in developmental programs coinciding with hatching has been observed in other systems such as *C. elegans* (Nicosia et al., 2013).

Based on the C4da developmental trajectory, we derive a morphological model containing two different growth programs. The inside-out program describes a growth type that is driven by tip branching and growth (Hannezo et al., 2017) while the outside-in program corresponds to more longer-ranging outgrowths followed by interstitial branching (Uçar et al., 2021). This model introduces a new spreading pattern parameter *sp* that determines the relative contribution of each growth type to development, which in turn determines the temporal dynamics of dendritic outgrowth while preserving other dendritic features. We were thus able to model a range of cell-type-specific differentiation trajectories for granule cells (*c*.*f*. Gonçalves et al., 2016; Radic et al., 2017, *in vivo* and *in vitro*) and for fly nociceptive neurons (Gerhard et al., 2017, *in vivo*). Our models also make predictions for structures and trajectories with temporally- or spatially-specific growth (which could be driven by changes in intrinsic or extrinsic growth cues) using multiple growth types. This includes the temporally distinct multi-phasic growth of C4da cells which we dissect here extensively; the spatially distinct short terminal branches (STBs) of class III cells, which have previously been shown to have different underlying microtubule structures compared to the rest of the cell (Stürner et al., 2022); and the branch compartmentalisation in pyramidal cells, where apical, basal, and oblique branch growths are differentially affected by different neurotrophins and other factors (Romand et al., 2011).

A key question arising from the observed developmental transitions is whether changes in growth programs disrupt wiring optimality. Despite the change in growth programs, C4da dendrites followed optimal wiring principles across all developmental stages, as shown using simple MST models (**Figure ??**, Nanda et al., 2018). Optimal wiring follows from optimal allocation of resources by balancing costs for total circuit wiring and signal propagation speeds (Chklovskii, 2004; Stepanyants and Chklovskii, 2005; Cuntz et al., 2007; Wen and Chklovskii, 2008; Ferreira Castro and Cardona, 2025). At the cellular level, such constraints have been shown to have several implications: (1) They lead to precise scaling relationships between measures of branching structure and connectivity, suggesting that different dendritic morphologies follow similar underlying growth rules (Cuntz et al., 2012); (2) They predict branching angle distributions (Kim et al., 2012; Bird and Cuntz, 2019); and (3), As shown in this work, they necessarily lead to optimal space filling regardless of target distribution, meaning that the dendritic span is optimally filled for a given optimal dendritic length (**Figure 4A, ??C**). Both scaling relationships and angle distributions were independent of the growth programs (**Figure 4**), and fitted the biological data throughout development.

Taken together, the distinct growth programs observed *in vivo* appear to preserve similar functional outcomes. This suggests a form of degeneracy in neuronal growth, whereby different developmental trajectories converge on comparable wiring solutions (Albantakis et al., 2024). Incidentally, the inside-out growth is reminiscent of the algorithmic iterations behind the greedy MST implementation of optimal wiring (Cuntz et al., 2007, 2010) (**Figure ??B**). The outside-in growth in addition is somewhat reminiscent of rapidly exploring random trees (rrt*) from graph theory which were previously linked to space filling (Kuffner and LaValle, 2011).

We have previously proposed our scaling equations from Cuntz et al. (2012) as a signature for optimal wiring. Importantly, recent work has shown that optimal wiring alone is not necessary for the power relationships in those equations (Ouyang et al., 2025, *i*.*e. L* ∼ *N* ^2/3^ in 3D or *L* ∼ *N* ^1/2^ in 2D). However, the particular scaling relations are a a consequence of the developmental implementation of optimal wiring and are not trivially achieved by stochastic growth. This contrast highlights that scaling laws cannot be interpreted independently of explicit growth models. Only through precise modelling can the relevant scaling relationships be identified, allowing fine-grained predictions of growth changes across development.

A limitation of the present study is that the growth rules and optimisation principles we identify are inferred from morphological dynamics rather than directly linked to underlying molecular or biophysical mechanisms. While our time-lapse imaging constrains the timing and nature of developmental transitions, the temporal resolution is insufficient to resolve the fine-scale dynamics of branch elongation, retraction, and local interactions that likely mediate these processes (Chalmers et al., 2016; He and Cline, 2011). In addition, our modelling framework is phenomenological and focuses on wiring and space-filling constraints, without explicitly incorporating intracellular signalling, cytoskeletal dynamics, or activity-dependent plasticity. Future work could address these limitations by linking model parameters to mutant phenotypes or elements of structural plasticity (Tavosanis, 2012; Yalgin et al., 2015), and by directly probing branch–branch repulsion and molecular cues that establish dendritic territories (Hattori et al., 2009; Matthews et al., 2007; Parrish et al., 2007).

Here, we provide a framework that links dendritic growth dynamics, space filling, and wiring optimality during development. Considerations of simultaneous multi-dendritic development and the spread of neurons into existing tissue would help make predictions about neuron packing (e.g. Braitenberg, 2001), neuron avoidance (Soba et al., 2007), gap filling (e.g. during repair, Song et al., 2012), and extracellular signal interactions, and would allow the exploration of how spatial constraints affect optimal wiring and space filling. Ultimately, our model links neurite outgrowth and space filling with circuit formation (Schrö ter et al., 2017; Helmstaedter, 2026) in neural networks of arbitrary complexity, enabling simulations based on well-understood principles rather than vast collections of neuronal reconstructions (Markram et al., 2015).

## Acknowledgments

We are grateful to M. Schölvinck, P. Soba, T. Stürner and A. Ziegler for comments on an earlier version of the manuscript and to C. Lossnitzer for helpful comments on the computer code. This work was supported by a BMBF grant (No. 01GQ1406 — Bernstein Award 2013), a DFG grant (CU 217/2-1), and by the German Federal Institute for Risk Assessment (Grant Agreement Number 60-0102-01.P636). The authors declare to have no competing financial interests.

## Author contributions

R.R., L.B., G.T. and H.C. designed the study. L.B. and G.T. designed and performed the experiments. R.R., A.F.C, and H.C. analysed the data, designed the growth model and performed all simulations. R.R., L.B., A.F.C, P.J., G.T., and H.C. wrote the paper.

## Methods

### Fly handling

To label Class IV da (C4da) neurons specifically, we used a fly strain that carried a fusion of a fraction of the enhancer of the *ppk* gene and eGFP (*w*^1118^;; *ppk* − *eGFP*) (Grueber et al., 2003). This strain expresses eGFP in C4da neurons from embryonic stage 16 throughout larval stages. We concentrated on the dorsal ddaC neuron (Grueber et al., 2002). For embryo collection, young fertilised females were placed in population cages, supplemented with apple agar plates maintained in an incubator at 25^*°*^*C* and 60% relative humidity. To obtain developmentally synchronised embryos the agar plate was supplemented with fresh yeast paste to promote laying of retained eggs and replaced after one hour. Females were then allowed to lay eggs for 30*min*. The collected eggs were placed back in the incubator and allowed to develop for 18*h*, when they were harvested, dechorionated with bleach for 3.5*min* and rinsed with water. Not all embryos were dechorionated by this gentle treatment, and the chorion was then removed mechanically with a brush.

### Time-lapse image acquisition

We developed a novel method to image the living larvae over several consecutive days causing as little interference as possible. Briefly, after each imaging session individuals were kept at 25^*°*^*C* and 60% relative humidity in a separate 500*µl* Eppendorf tube filled with 200*µl* standard cornmeal-agar fly food. To regain the test subject the fly food was dissolved in water and the larvae were localised under a binocular microscope (**Figure 1A**). To keep animals immobilised for imaging, but alive through the imaging sessions, we utilised three different physical immobilisation techniques at different stages. **Embryo**: We created a grid with 6 chambers of approximately 1*mm* × 0.5*mm* × 0.5*mm* on an object slide using adhesive strip. Dechorionated embryos were placed in the grid, covered with halocarbon oil 700 (Sigma) and oriented in such a way that the dorsal class IV ddaC neurons faced towards the coverslip. **L1 and L2 larvae**: Images of L1 and L2 larvae were acquired under a custom-made object slide (Dimitrova et al., 2008). The metal custom object slide fitted a round coverslip onto which the larva was placed, oriented and covered with halocarbon oil. A plastic net on a round plastic support was then placed on top of the coverslip and fixed with screws to the metal slide. Detailed information about the design is available upon request. **L3 larvae**: L3 larvae were physically immobilised in halocarbon oil in-between an objective slide and a coverslip. Double-sided adhesive tape served as spacer. On average two out of five individuals survived through all imaging steps. During this procedure, development until the pupal stage was extended from typical five days at 25^*°*^*C* to about seven days in our preparation. Given the increased development time in these conditions from ∼ 120*h* to ∼ 168*h*, embryo images were taken 18*h* − 19*h AEL*, L1 images at 42*h AEL*, L2 at 90*h AEL* and L3 images 138*h AEL*. The measured ddaC dendrite size at 90*h* and at 138*h* corresponded to our previous measures of L2 and feeding L3 animals in control conditions, respectively. Images were acquired with a Leica TCS SP2 Confocal Microscope (http://www.leica-microsystems.com), maintaining minimal laser intensity to reduce tissue damage. To avoid issues due to larval movement, we minimised imaging time per *z*−stack and thus chose a relatively large *z*−step of 2*µm* combined with a larger pinhole size (85− 150*µm*) to obtain a voxel depth of 1.7− 2.2*µm*. For embryos, we used a 40× oil immersion objective and for all the other stages a 20× oil immersion objective.

### Data analysis

All data analysis was performed in *Matlab* (www.mathworks.com) using our own software package, the *TREES toolbox* (www.treestoolbox.org, version 1.15 and interim version, Cuntz et al., 2010, 2011). A number of new TREES toolbox functions were custom-made for this study and will be incorporated in the existing *TREES toolbox* with the publication of this work: span tree, scaleS tree, scaleV tree, theta tree, compare tree, ui comp tree, theta mc tree, growth tree. See below for details on the individual functions. In the following, function names are in monospaced font with *TREES toolbox* functions having a tree suffix. All data will be made available on www.NeuroMorpho.Org and all new *TREES toolbox* functions will be made available on www.treestoolbox.org. The entire code and data will be archived online on Zenodo.

### Anatomical data of C4da neurons

Image stacks from the confocal microscope were imported in the *TREES toolbox* and manual reconstructions of all dendrites were performed individually (total amount of neurons *N* = 168) using the dedicated reconstruction user interface cgui tree. During the reconstruction process we determined (1) adequate internode distances i.e. spatial resolution at which to resample (resample_tree) the dendritic structures, (2) parameters for a diameter taper toward the root (quaddiameter tree) that best reproduced the overall diameter values of the real cells, (3) soma length and soma diameter values to best map soma diameters (soma_tree) *post hoc* on the existing reconstructions, and (4) region codes for dendrite, soma and axon. Axon nodes were discarded for the remaining analysis.

### Classical branching statistics

A large palette of branching statistics were collected for each dendrite reconstruction separately using simple combinations of *TREES toolbox* functions: (1) **Total dendrite length** *L* was the sum of all internode lengths (len_tree), (2) **number of branch points** *B* were calculated as the sum of all nodes that are branch points (B_tree), (3) covered **surface area** *S* was calculated with a novel custom written *TREES toolbox* function span_tree that counts 1 − *bins* after morphological closing with a disc of 4*µm* radius of a binary matrix with 1 − *bins* at tree node locations after resampling the trees to 1*µm* internode distances – similarly to previous work on fly lobula plate tangential cells (Cuntz et al., 2008), (4) **Sholl analysis** (Bird and Cuntz, 2019) that calculates the number of intersections of a tree with a growing circle centered on the dendrite root was done by using the dedicated *TREES toolbox* function sholl_tree, (5) normalised Sholl analysis was done by normalising the Sholl radii to 6× the mean Euclidean distance in the tree (eucl tree) and normalising the number of intersections with the integral over the Sholl intersections diagram, (6) **branch order** distributions were calculated by identifying how many nodes and the overall dendritic cable length of their connected segments in the tree had a given branch order (BO_tree), (7) normalised branch orders were normalised to the next integer of 2.5× the mean branch order in any given tree and the cable distribution was normalised to the total length of dendrite, (8) **branch length** distributions were obtained by combining two *TREES toolbox* functions that calculate the path length from each node to the root (Pvec_tree) and that dissect the trees into their individual branches (dissect_tree), (9) normalised branch lengths were normalised to 3× the mean branch length in the tree and branch length frequency was normalised to the integral over the distribution, (10) **compression** value distributions were calculated as the ratio between Euclidean distances (eucl_tree) and the path distances (Pvec_tree) to the root for all nodes, (11) the **Strahler order** distributions were calculated as the percentages of dendrite length per given Strahler order (strahler_tree) (Vormberg et al., 2017), (12) **branch angle** distributions were obtained by calculating the angles in the branching plane of each branch point (angleB_tree). Overall, we chose not to normalise to the maximum value but rather to a multiple of the mean because maximum values are more sensitive to noise.

### Scaling of real trees

In order to compare real trees with a simple maximal space filling example, all reconstructions were scaled to the same surface area *S* as a reference diamond (*h* · 1, 000, 000*µm*^2^, see supplementary **Figure ??** and supplementary section **‘Mathematical calculations for maximal space filling’)** by scaling the trees twice while measuring *S* (span_tree, see above) to make sure that target *S* is reached precisely in all cases (new *TREES toolbox* function scaleS_tree). Real trees in **Figures 2D-F, 5**, **supplementary ??** were scaled in such a manner.

### Space-filling measure 1*/θ*

A custom-written new *TREES toolbox* function theta tree was used to calculate how space filling a tree is. *θ* is the distance one has to move away from the tree to cover 90.69% of the spanning surface of the tree, in analogy to our maximal space filling mathematical framework derived from packing circles, see **Supplementary text**. Similar to the surface area *S* calculation, we calculated a binary matrix with 1 − *bins* after morphological closing with a disc of 4*µm* radius and used the *Matlab* function bwdist to calculate the distances from each bin within this spanning area to the closest bin occupied by the dendritic tree. Similar ‘box-counting’ methods have appeared in the literature (Wen et al., 2009; Shree et al., 2022), but the approach here allows the calculation of both single values of *θ* and the entire detailed distance distributions in histograms. The iterative growth process followed a slightly different but equivalent approach using a Monte Carlo method since for 3D cases the binary matrices would have been very large (see section **‘Iterative growth model**’).

### Time-lapse analysis

Before registration of the branch and termination points, the morphologies of identified pairs of dendrites (the same cell in two consecutive time steps, *N* = 47 pairs) were rotated, translated and scaled separately in both dimensions to maximise the overlap of dendritic branches. Branch and termination points were then registered first manually and second using a fully-automated function (new *TREES toolbox* function compare_tree, see **Figure ??**).

For the stretching analysis, the individual scaling steps were ignored and the vectors were drawn between branch and termination points of time points 1 and 2 for all pairs.

We took advantage of both manual and automated of registration sets to track and and quantify the branching changes across the structures. Considering the branches formed between the soma and the tracked branching and termination points to be common between the structures in a given pair, we were able to isolate retracted branches (branches that appear in structure 1 and not in 2) and newly formed branches (branches that appear in structure 2 and not 1 and start from non-termination points). Branch elongation and shrinkage processes were identified as (1) the growth of untracked branches (similar to newly-found branches) from termination points and (2) the movement of termination points tracked with compare_tree from one time point to the next (since the fully-automated method stringently followed termination points, see **Figure ??**E).

### Dendritic minimum spanning trees

Optimal wiring was implemented as previously described (Cuntz et al., 2007) with a minimum spanning tree algorithm connecting a given set of targets to minimise (1) total cable length and (2) path lengths from any target along the tree towards the root of the tree. This second cost was weighted by the balancing factor *bf*, the only parameter that the wiring algorithm takes: *wiring cost* = *L* + *bf* · *PL*. This method was successfully used previously (Cuntz et al., 2008, 2010) and all algorithms are available in the *TREES toolbox* (www.treestoolbox.org,MST_tree).

Targets were distributed uniformly in the surface area of each real dendrite, one at a time (**Figure ??A**), and connected them with the minimum spanning tree algorithm. We generated minimum spanning trees connecting as many targets as required to reproduce the number of branch points *B* in the original tree and matched the resulting synthetic tree surface area *S* each using the same scaling procedure used for real reconstructions (see above). We then scanned the parameter *bf* of the model between 0 and 1 in steps of 0.025. An error function comparing mean total length, branch order, compression and path length between synthetic trees and their real counterparts yielded an optimal *bf* of 0.225. The corresponding synthetic trees were then used with this *bf* value (**Figure 4D**). For visual purposes, all synthetic trees were smoothed (smooth_tree) and a slight jitter was added (jitter_tree) that did not affect the branching statistics.

### Iterative growth model

The iterative growth model (new TREES function growth_tree) is an extension of the original minimum spanning tree model implementing optimal wiring. It was fit to cover the two developmental programs uncovered from reconstructions of dendrite development, and the span of developmental trajectories between them. Given a surface area or volume in which to grow a dendrite, this span is probed with randomly placed target points, with every iteration of the algorithm growing towards one of these points to simulate a sequential growth process. The number of target points is chosen by finding the lowest number that allows the algorithm to reach the set termination condition before it runs out of points to connect to. This is done by starting with (roughly) twice the number of branching points in the reference structure, and decreasing that number until the algorithm stops prematurely. Selecting the target point to grow to and where to connect it on the tree is based on optimising two separate functions: minimising wiring cost, as described in the previous section, and maximising space filling, which is based on how far a given target point is from the existing tree structure. These two optimisation procedures are combined in a single scoring function used to select the most optimal tree node to target point combination in a given iteration: *score* = ((1 − *sp*) ∗ *wiring cost*) − (*sp* ∗ *node distance*), where the scores pertaining to each function are normalised to 1 and balanced based on the value of the spreading pattern (*sp*) parameter. At *sp* = 0, the algorithm prioritises connections that minimise the wiring cost, while at *sp* = 1 the algorithm connects preferentially to points that are further away from the tree (while still minimising wiring costs as much as possible). An additional parameter *k*, which is internally saturated using *tanh*(4*k*), adds uniform random noise to the scoring function. Following these parameters, the tree node – target point combination with the lowest calculated score is chosen and added to the tree. The new branch is then subdivided into 1*µm* segments and jittered slightly before the next iteration starts. The growth process stops when a termination condition is reached, which can either be running out of target points or stopping when a target total length *L* or number of termination points *T* is reached.

### Synthesised morphologies of other planar and 3D dendrites

Five synthesised dendrites were grown from scratch using the growth tree iterative growth model. Synthesised **Fly lobula plate tangential cells** (Cuntz et al., 2008) **were obtained using** *bf* = 0.6, *sp* = 0.8, *k* = 0.4, *radius* = 800*µm*; **Rat dentate gyrus granule cells** (Rihn and Claiborne, 1990) were obtained using *bf* = 0.975, *sp* = 0.3, *k* = 0, *radius* = 400*µm*; **Rat cortical layer 5 pyramidal cells** (Wittner and Miles, 2007) were obtained using *bf* = 0.9, *sp* = 1 *k* = 0.5, *radius* = 1, 000*µm*; **Fly CIII dendritic arborisation sensory neurons** (Wittner and Miles, 2007) were obtained using *bf* = 0.9, *sp* = 1 *k* = 0.5, *radius* = 1, 000*µm*.

### Intermediate structure prediction

To validate our synthesised developmental trajectories, we tested our model’s ability to predict the correct intermediate structures by comparing our model’s iterations to earlier phases of real neurons. Two electron microscopy datasets containing overlapping neurons were used here: one from the fly L1 stage taken from the A1 segment, and another from the L3 stage/A3 segment. Six neuron types were isolated (A10a, A09a, A09c, A09l, A02m, A02n) from both datasets, and preprocessed by separating the dendrites, axons, and somatic linker cables. This paradigm then involved modeling each target L3 dendrite and combing through the iterations leading to it to find the L1-equivalent structures, which were defined as those closest length-wise to the real L1 dendrites.

Similar parameters were used to generate the structures for all six cell types: **A10a neurons** were obtained using *bf* = 0.2, *sp* = 0, *k* = 0.5, *radius* = 80*µm*; **A09a neurons** were obtained using *bf* = 0.1, *sp* = 0, *k* = 0.6, *radius* = 80*µm*; **A09c neurons** were obtained using *bf* = 0.1, *sp* = 0.1, *k* = 0.6, *radius* = 80*µm*; **A09l neurons** were obtained using *bf* = 0.25, *sp* = 1, *k* = 0.6, *radius* = 200*µm*; **A02m neurons** were obtained using *bf* = 0.1, *sp* = 0, *k* = 0.5, *radius* = 120*µm*; **A02n neurons** were obtained using *bf* = 0.05, *sp* = 0.2, *k* = 0.6, *radius* = 80*µm*.

The connectome model graphics in **Figure 6B** were generated by concatenating the modeled dendritic structures to the previously discarded axon and linker cable segments. In order to somewhat account for tissue stretching between the two stages, L1 structures were scaled down according to the overall difference in size between the real connectomes (using the horizontally protruding somata as reference).

