## Supplemental material for "A developmental stretch-and-fill process that optimises dendritic wiring"

### Supporting information

#### Validation of the manual registration using an automated method

In order to check the robustness of our time lapse analysis results of C4da growth (**Figure 2**), we also performed the point registration across structures using a fully automated method (Hermila et al., 2026), the `compare_tree` function (**Figure S1**). Repeating the stretching analysis shown in **Figure 2** panels B and C showed similar results between the registrations (**Figure S1B, C**), down to the very similar slope values for stretching amplitude vs distance from the soma.

There were some differences in the registrations themselves: we found that 54% of all node registrations were concurrent, where both methods performed the same registrations; 22.5% of the registrations were differing, where each method assigned nodes from one structure to different nodes in the next; and the remaining 23.5% of registrations were separately matched, which included the nodes that were only assigned by one method and not the other (**Figure S1D**).

We were able to isolate two main differing registration patterns: misaligned tips and missing branches (**Figure S1E**). In the first case, `compare_tree` consistently assigned termination points in one structure to termination points in the next, even when the same points were assigned to continuation points in the manual registrations. In most cases, this was a matter of tip growth – the `compare_tree` function followed the tip to its new position while manual registration did not and considered the previous location of the tip (which is now no longer a tip) to be the corresponding point in the following structure. There is no strictly correct or incorrect version here, as this depends on how the moving tip is defined. In the second case, `compare_tree` sometimes missed entire branches between the structures, with up to 76 nodes in a single missed cluster.

Overall, the analysis exemplifies that both automated and manual registration methods are useful. In our case, the automated registration corroborates our findings from manual registration. The latter are more detailed but are also more likely to be prone to subjective biases.

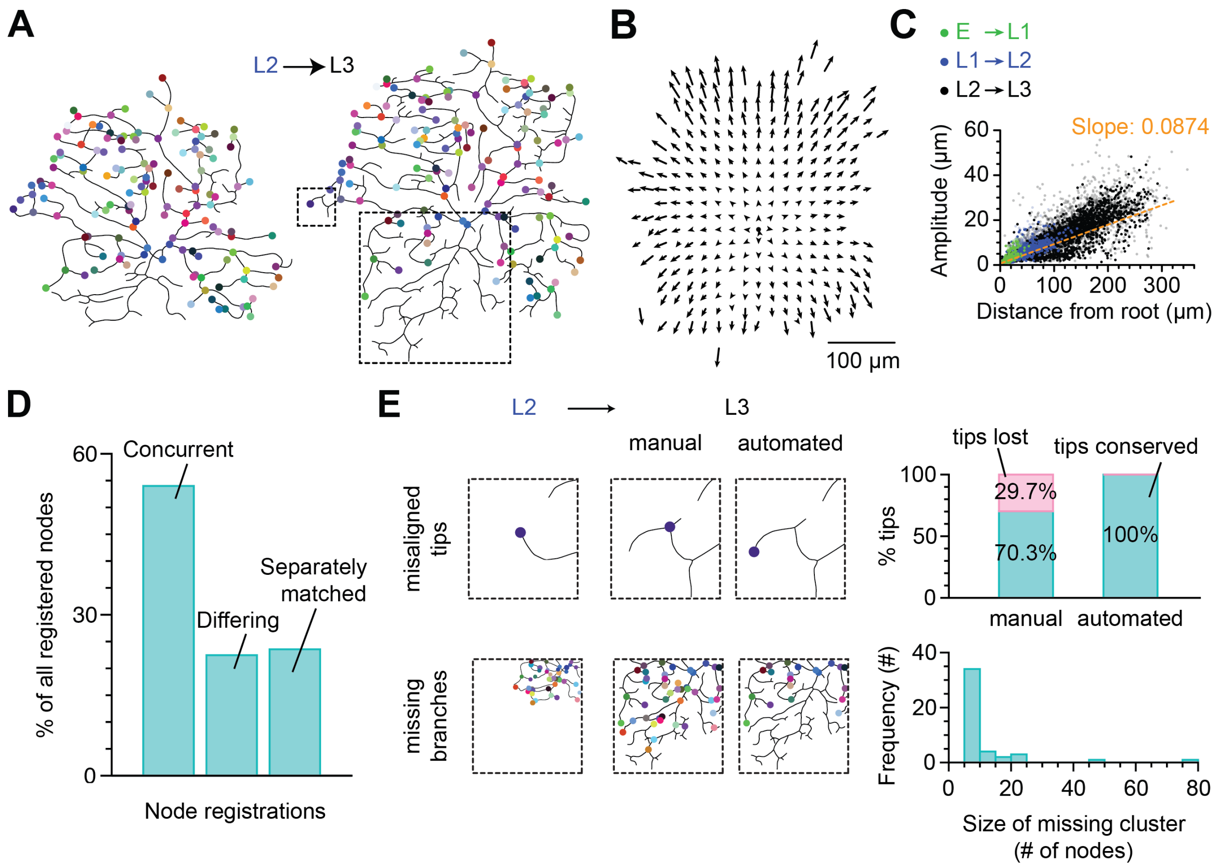


**Figure S1. Comparison of manual and automated time point registrations.**

**A**, Branch and termination points (coloured dots) in time 1 (left, here L2), are registered to corresponding points in time 2 (right, here L3, same colours). **B**, Same analysis as in **Figure 2B**: overall stretching analysis for all pairs ($N = 47$). Arrows (every $25\mu m$) are averages only shown where data for at least 5 pairs were available. Dendrites were centred on their root (black dot). **C**, Same analysis as in **Figure 2C**: Analysis of stretching by comparing the arrow lengths collected from all pairs). Greyed out points represent tips, linear fit (dashed orange line) determines the slope. **D**, Left, diagrams showing examples of the main sources of differences between the manual and automated registration methods: misaligned tips (top row, small dashed rectangle in **A**) and entire missing branches (bottom row, larger dashed rectangle in **A**). Right, graphs quantifying each observed difference. **E** Bar plots quantifying in how many samples a given percentage of the nodes was concurrently registered between the two methods (top, green), differently registered (middle, gray), and registered in one method but not the other (bottom, pink).

#### Optimal space filling measure from hexagonal lattice considerations

Efficient coverage of dendrites on a given surface area could be achieved by spreading out branches to regularly distributed target locations. As a reference, we used targets on a closed hexagonal lattice arrangement that are well spread (**Figure S2A**) (Hales, 2000). Tightly packed circles with radius $\theta$ around these targets are known to cover $\frac{\pi}{2\sqrt{3}} \approx 90.69\%$ of total available $S$. This simple geometrical framework yields direct relationships for all morphological variables of interest when connecting $N$ such targets to a tree:

$$\theta = (h^{-1/2}/2) \cdot S^{1/2} \cdot N^{-1/2} \approx \frac{S}{(2 \cdot h \cdot L)} \qquad L \approx h^{-1/2} \cdot S^{1/2} \cdot N^{1/2} \tag{1}$$

where $h = \frac{\sqrt{3}}{2}$ (**Methods**). In order to compare these relations with random target distributions (**Figure S2B**) and real trees (**Figure S2C**), one can calculate their corresponding equivalent $\theta$ values as the distance one has to move away from all points on the tree to cover $\sim 90.69\%$ of $S$. Equipped with **Equation 1** and after normalising $S$ in all cases we were able to isolate entirely linear relationships between the square root of the number of target points $N^{1/2}$, $L$ and $\frac{1}{\theta}$, the inverse of $\theta$ quantifying the amount of space filling (Stepanyants et al., 2002).

Iterating over $n$ and starting at $n = 2$ we distribute $N = n^2$ targets on a hexagonal grid within diamond-shaped boundaries, $n$ being the number of targets on an edge of the diamond (**Figure S2A**). The targets form a space divided into equilateral triangles where neighbors are distance $l$ away from each other. If the basis of one triangle is set to be horizontal, the height of the equilateral triangle relates to $l$ with $h = \frac{\sqrt{3}}{2}$ and the total surface of the diamond is:

$$S = h \cdot (n-1)^2 \cdot l^2. \tag{2}$$

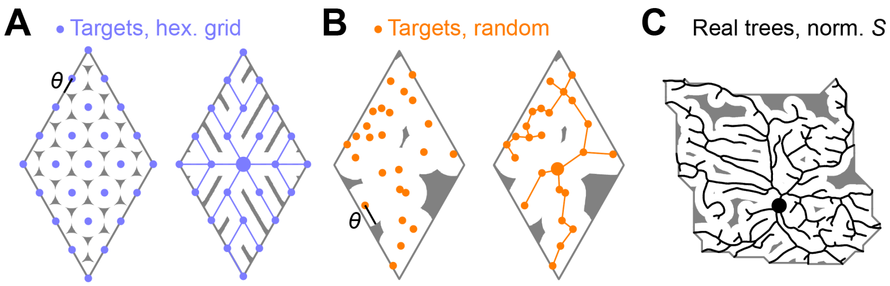


**Figure S2. Optimal space filling on a hexagonal lattice.**
**A**, Targets (blue dots) distributed on a closed regular hexagonal grid optimise space coverage (*left*) and form a minimum spanning tree (blue straight lines) when connected to minimise total cable and path lengths toward the root (large blue dot, *right*). Grey shaded area indicates $\sim 10\%$ ($> \theta$; $\theta$ shown as black line for one sample target) furthest away from targets (*left*) or from the tree (*right*). **B**, Similar arrangement as **A** but with random uniformly distributed targets (orange colours) as a comparison. **C**, Similar arrangement as **A** for a real C4da dendrite normalised to the same surface area $S$ as in **A** (targets are not shown since they are unknown).

Circles of radius $\theta = \frac{l}{2}$ surrounding the targets on the hexagonal grid are packed optimally and fill the surface area. It can easily be shown that the surface area covered by the circles within the diamond shape follows $S' = (n-1)^2 \times \textit{the surface of the circle}$ since the four corners together cover the surface of one circle, each edge point covers half of a circle and each center point covers one entire circle each, so:

$$S' = \frac{\pi}{4}(n-1)^2 \cdot l^2, \tag{3}$$

which corresponds to a portion of $\frac{\pi}{2\sqrt{3}} \approx 90.69\%$ of the total surface $S$ (see **Equation 2**).

Combining **Equation 2** and $\theta = \frac{l}{2}$, we find that $\theta$ decreases with $N$ and increases with $S$ in

$$\theta = \frac{h^{-\frac{1}{2}}}{2} \cdot S^{\frac{1}{2}} \cdot N^{-\frac{1}{2}} \tag{4}$$

Connecting the points on the grid optimally (in terms of total length of cable) yields a tree with length $L = (N-1) \cdot l$ since for each target (except the first) exactly one piece of length $l$ is required to connect it to the rest of the tree. Note that the minimum spanning tree has many solutions since all the distances between neighbors are equal, being precisely $l$. An inherent relation exists between $L$ and $\theta$:

$$L = 2\,(N-1) \cdot \theta \tag{5}$$

Combining $L = (N-1) \cdot l$ with $N = n^2$ and **Equation 2** results in

$$L = h^{-\frac{1}{2}} \cdot S^{\frac{1}{2}} \cdot \frac{(n^2-1)}{(n-1)} \approx h^{-\frac{1}{2}} \cdot S^{\frac{1}{2}} \cdot N^{\frac{1}{2}} \tag{6}$$

reminiscent of the power law that we described previously for planar dendritic minimum spanning trees generally (Cuntz et al., 2012) and in this case providing an upper bound. Further combining **Equations 6 and 4**:

$$\theta \approx \frac{S}{2 \cdot h \cdot L} \tag{7}$$

For a $2D$ square grid, the equations are similar although the coverage $\frac{S'}{S} = \frac{\pi}{4} \approx 78.54\%$ is not as good with:

$$S = (n-1)^2 \cdot l^2 \qquad \theta = \frac{1}{2} \cdot S^{\frac{1}{2}} \cdot N^{-\frac{1}{2}} \qquad L \approx S^{\frac{1}{2}} \cdot N^{\frac{1}{2}} \qquad \theta \approx \frac{S}{2 \cdot L}$$

while $N = n^2$, $\theta = \frac{l}{2}$, and $S' = \frac{\pi}{4}(n-1)^2 \cdot l^2$ remain unchanged.

Similarly, a $3D$ cubic grid can be constructed with:

$$V = (n-1)^3 \cdot l^3 \qquad \theta = \frac{1}{8} \cdot V^{\frac{1}{3}} \cdot N^{-\frac{1}{3}} \qquad L \approx V^{\frac{1}{3}} \cdot N^{\frac{2}{3}} \qquad \theta \approx \frac{1}{8} \cdot V^{\frac{1}{2}} \cdot L^{-\frac{1}{2}}$$

and $N = n^3$, $\theta = \frac{l}{2}$, $V' = \frac{\pi}{6}(n-1)^3 \cdot l^3$ and a poorer coverage even of $\frac{V'}{V} = \frac{\pi}{6} \approx 52.36\%$ and a $\frac{2}{3}$ power between $N$ and $L$ as we have shown previously for $3D$ cases (Cuntz et al., 2012).

#### MST-based model reproducing C4da development

We checked that the C4da reconstructions satisfied optimal wiring criteria according to our minimum spanning tree (MST) based model (Cuntz et al., 2010). A good match was found for $bf = 0.225$ (**Figure S3**). The model then readily explained the scaling relationships in (**Figure 1**).

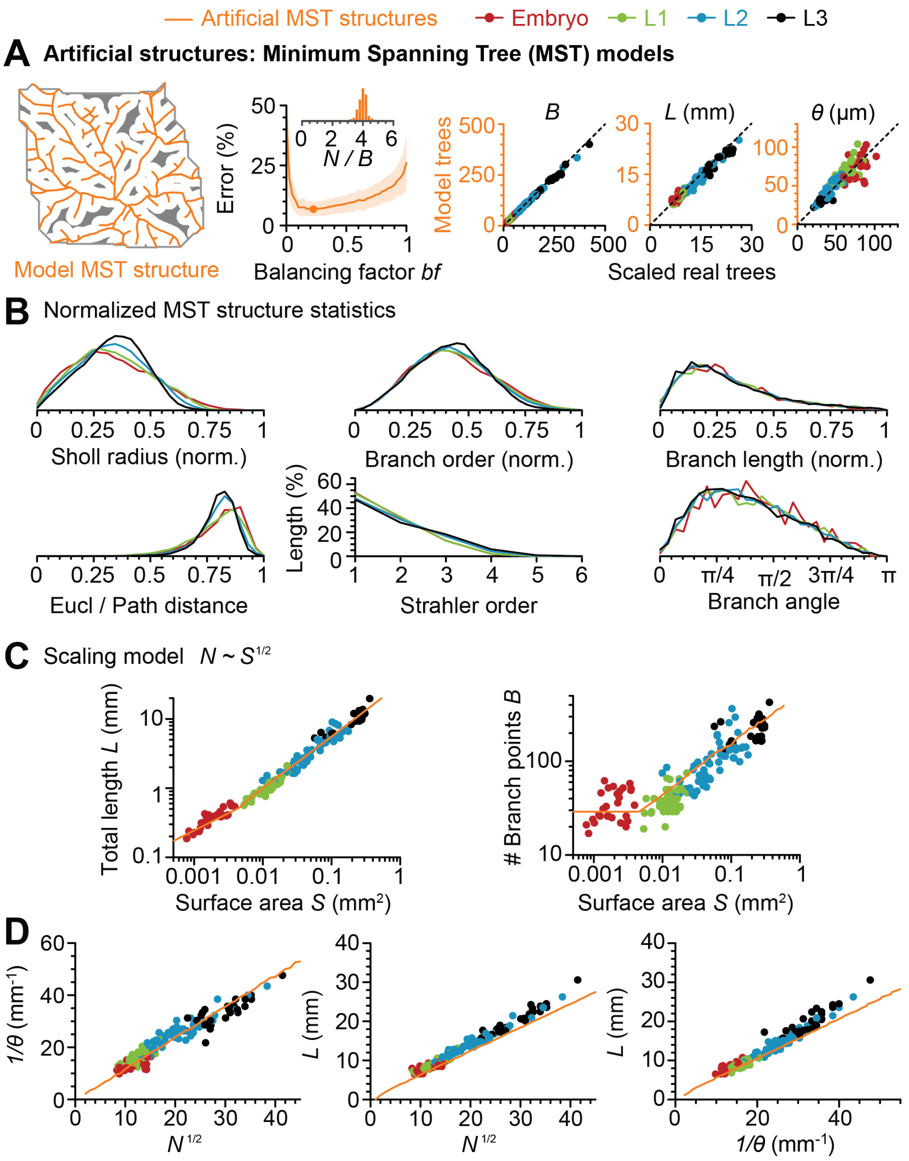


**Figure S3. MST models of C4da structures.**

**A**, Sample model trees based on homogeneously distributed targets in the dendrite spanning area (see **Methods**). *Upper row, left to right*: Example MST structure; (See next page.)

**Figure S3. (continued)** Model parameter $bf$ vs. error (comparing total length, branch order, compression and path length with corresponding real tree values; orange line), optimal $bf$ ($0.225$, orange dot), standard deviation of the error (shaded area), and distribution of ratio values between numbers of targets and branch points (inset, $4.07 \pm 0.29$) for the optimal $bf$; Validation of number of branch points $B$, total cable length $L$, and space coverage $\theta$ between scaled real trees and model trees with dashed line indicating unity for comparison. **B**, similar panels as in **Figure 1E** but for synthetic trees. **C**, Scaling behaviour of the MST model from **(A)** (orange) compared with data from the real structures (c.f **Figure 1F**). **D**, Comparison of the scaling behaviour of the variables in the space filling equations between real neurons (dots, same colours as in **Figure 1**) and the MST model (orange lines) shown in A. Number of branch points in real dendrites were converted to target number $N$ by the factor $4.07$ obtained in **A**, second panel inset.

#### Algorithm comparison

We compare here our new model with two other algorithms: the minimum spanning tree (MST) algorithm (Cuntz et al., 2010) and a growth function published in a previous version of this paper (bioRxiv version 1). **Figure S4** provides a comparison of the 3 algorithms, with panel **A** showing the structures generated by each algorithm throughout its parameter space. Most importantly, while all three have a balancing factor ($bf$) parameter that controls the wiring cost optimisation, the new algorithm presented in this version of the manuscript adds the $sp$ parameter that controls the dendritic spreading pattern during the growth of the structure. Roughly, $sp$ affects only the timing of development while $bf$ affects the final structures. Looking at the structures, this is not entirely the case, and is an issue that is exacerbated by the fact that the structures presented here are generated without any noise or jitter, making high $sp$ structures look particularly unrealistic. In order to see the full effect of the $sp$ parameter, we look at a time-lapse analysis of these structures. **Figure S4B** shows the intermediate structures iteratively generated by each algorithm leading up to structures with $bf = 0.5$ in **Figure S4A**. The developmental time-lapse of the MST structure (*top row*) shows a growth pattern reminiscent of the inside-out growth described in **Figure 4A**, while that of the original growth function (*bottom row*) shows a growth pattern similar to the outside-in growth type. The new algorithm (middle row) combines both types: as the value of the $sp$ parameter increases, the spreading pattern gradually changes from the inside-out growth at $sp = 0$ to outside-in at $sp = 1$.

Due to the nature of the optimisation carried out by the new algorithm, despite the fact that $sp$ changes linearly from 0 to 1, its effect is not linear or smooth. This can be seen in both the developmental profiles of the structures as an abrupt change in growth type around $sp = 0.5$ (**Figure S4B**) and in the quantification of these growth trajectories (**Figure S4C**), which look at how much of the growth is actually due to tip growth (*left*) and how the structures fill the entire dendritic span available to them (*right*) across development.

Note that despite the fact that the output of the new growth model at $sp = 0$ seems very similar to the output of the MST model, it is not identical, as the former allows inter-nodal interstitial branching while the latter does not.

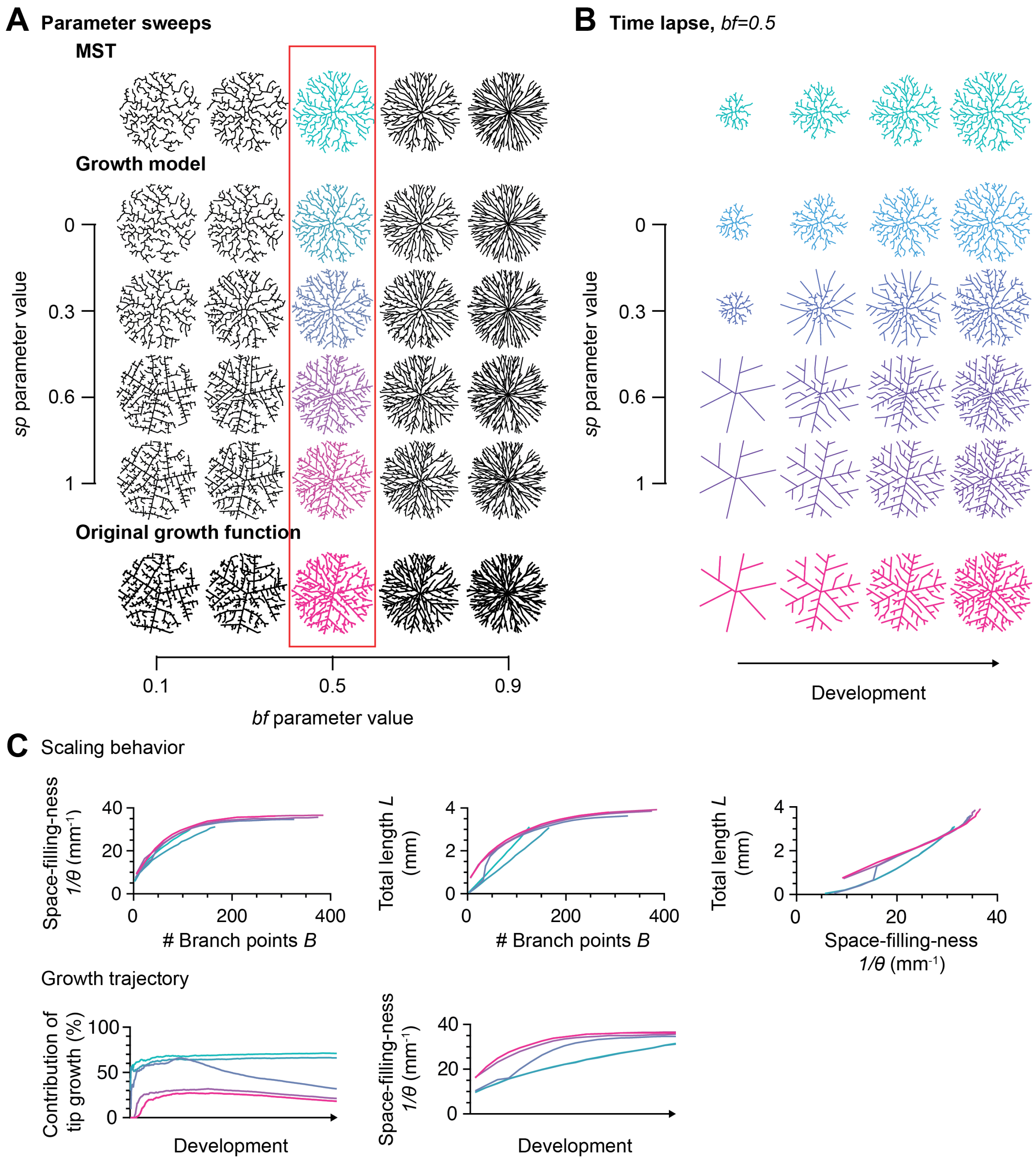


**Figure S4. Comparing algorithms of morphological models.**

**A**, Structures resulting from a parameter sweep of all 3 functions (MST, top row; new growth model, middle set of rows; original growth function, bottom row) for the *bf* and *sp* (new growth model only) parameters. **B**, Iterative growth of the coloured structures shown in the red rectangle in **A**. Structures across the rows are matched by total dendritic length, not developmental time. **C**, *Top row*: graphs showing the scaling behaviour of the total length, space-filling-ness, and number of branch points parameters to each other throughout development. *Bottom row*: Quantifying the growth trajectories of each structure by looking at the percent contribution of tip growth to the overall growth and at the space-filling-ness. The colours in the entire figure correspond to the algorithm used and/or the *sp* value.

#### The effect of stochasticity

We look here at the effect of increasing noise (the $k$ parameter) on the synthesised structures and their emergent developmental profiles. Beyond the obvious effect of noise making the structures look more realistic by breaking up the very straight lines (**Figure S5A**), it also interfered with the effect of $sp$ on the timing of growth during development (**Figure S5B**). Increasing stochasticity lead to the developmental trajectories imposed by different values of $sp$ becoming less distinct and eventually overlapping at $k = 1$ (**Figure S5C**, *left*). The amount of deviation from optimality varied for each $sp$ value but consistently increased with increasing $k$ (**Figure S5C**, *right*).

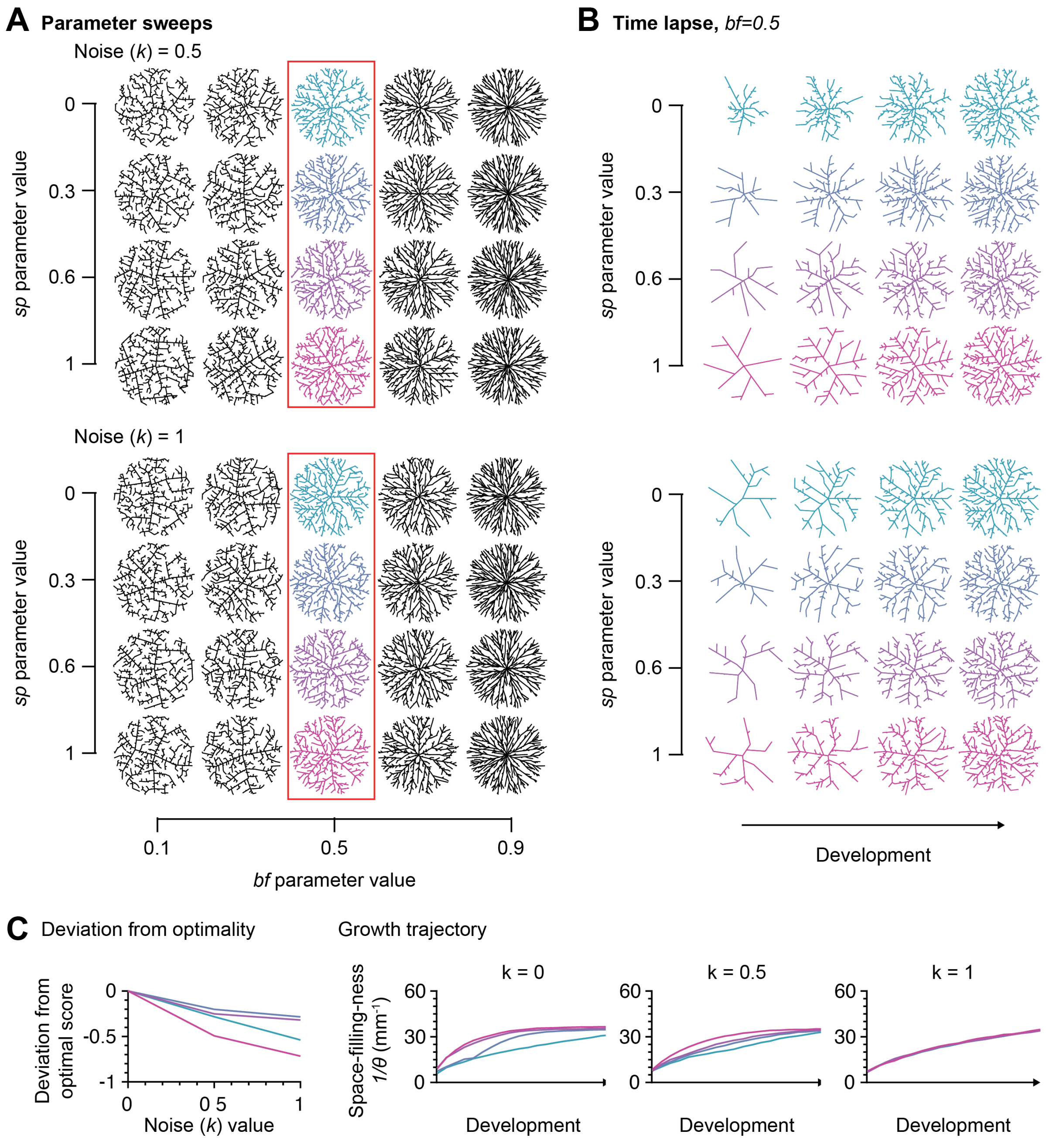


**Figure S5. The effect of stochasticity on the growth model.**

**A**, Structures resulting from a $bf$ vs $sp$ parameter sweep for new growth model with different values of the noise parameter $k$ ($k = 0.5$ top row, $k = 1$ bottom row). **B**, Iterative growth of the coloured structures shown in the red rectangles in **A**. Structures across the rows are matched by total dendritic length, not developmental time. **C**, Graphs quantifying the effect of $k$ on development: on the left, a graph showing how far from optimality the development deviates for each value of k; on the right, graphs showing how the evolution of space filling ness across development changes for each combination of $sp$ and $k$ values.

#### Single-step C4da development

In order to investigate the importance of the two-step developmental process of C4da cells (reproduced in **Figure S6A, B**), we simulated full C4da development with either the first or second growth step (**Figure S6C**).

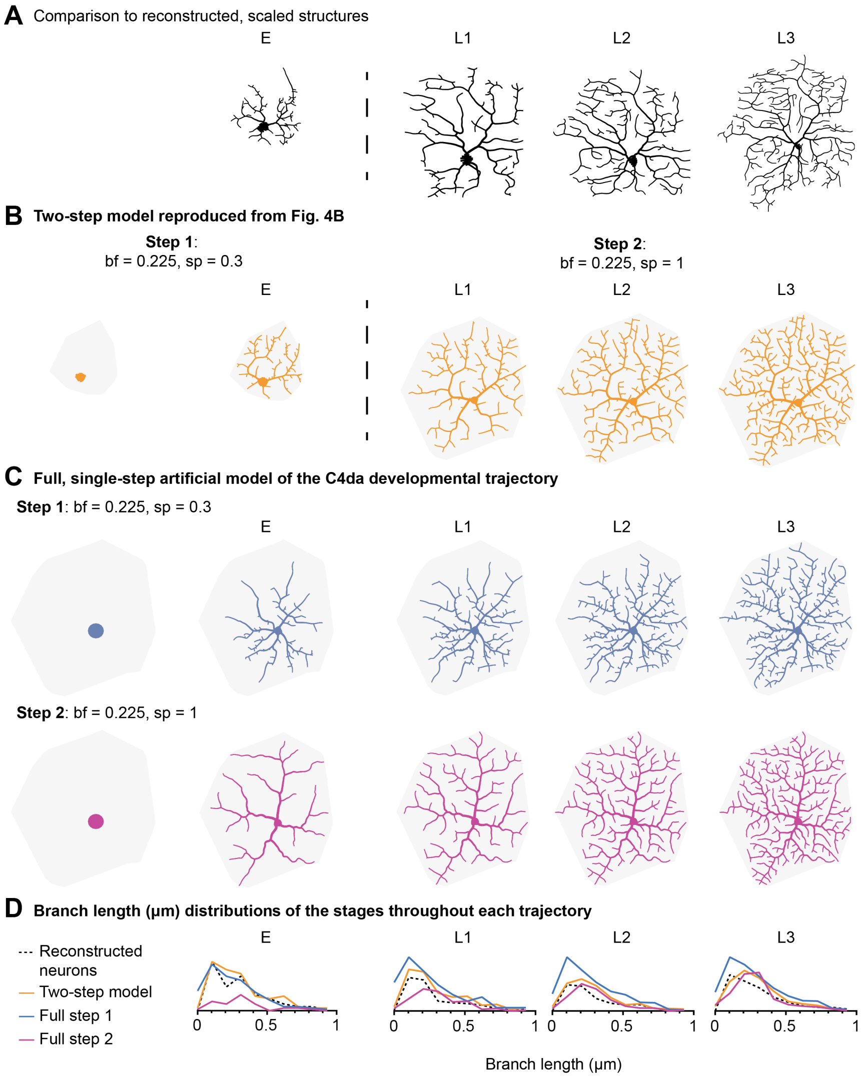


**Figure S6. Potential C4da developmental trajectories.**

**A, B**, Reproduced from **Figure 5A** for better comparison: **A**, Reconstructed C4da structures scaled appropriately, **B** the two-step model of C4da development. **C**, Alternative, single-step models of development, either growing entirely with the properties of the first step (*top*) or the second step (*bottom*). **D**, Distributions of branch length values in $\mu m$ at each stage for the reconstructed structures (*black*, dashed line) along with the synthesised two step (*orange*) or single step (*blue*, full step 1 trajectory; *purple*, full step 2 trajectory) structures.

In addition to the resulting structures deviating visually from both reconstructed and two-step model structures, the branch length distributions of each matched stage across all four trajectories also showed significant deviations (**Figure S6D**). While the full step 1 model matched the two-step model and the reconstructed data at the early stages, it deviated from them after the E stage; the full step 2 model on the other hand showed deviations at early but not later stages. This lines up with our prediction that step 1, inside-out-type growth is needed to reproduce early C4da structures while step 2 or outside-in-type growth is needed for the later ones.

#### Statistics for additional cell types

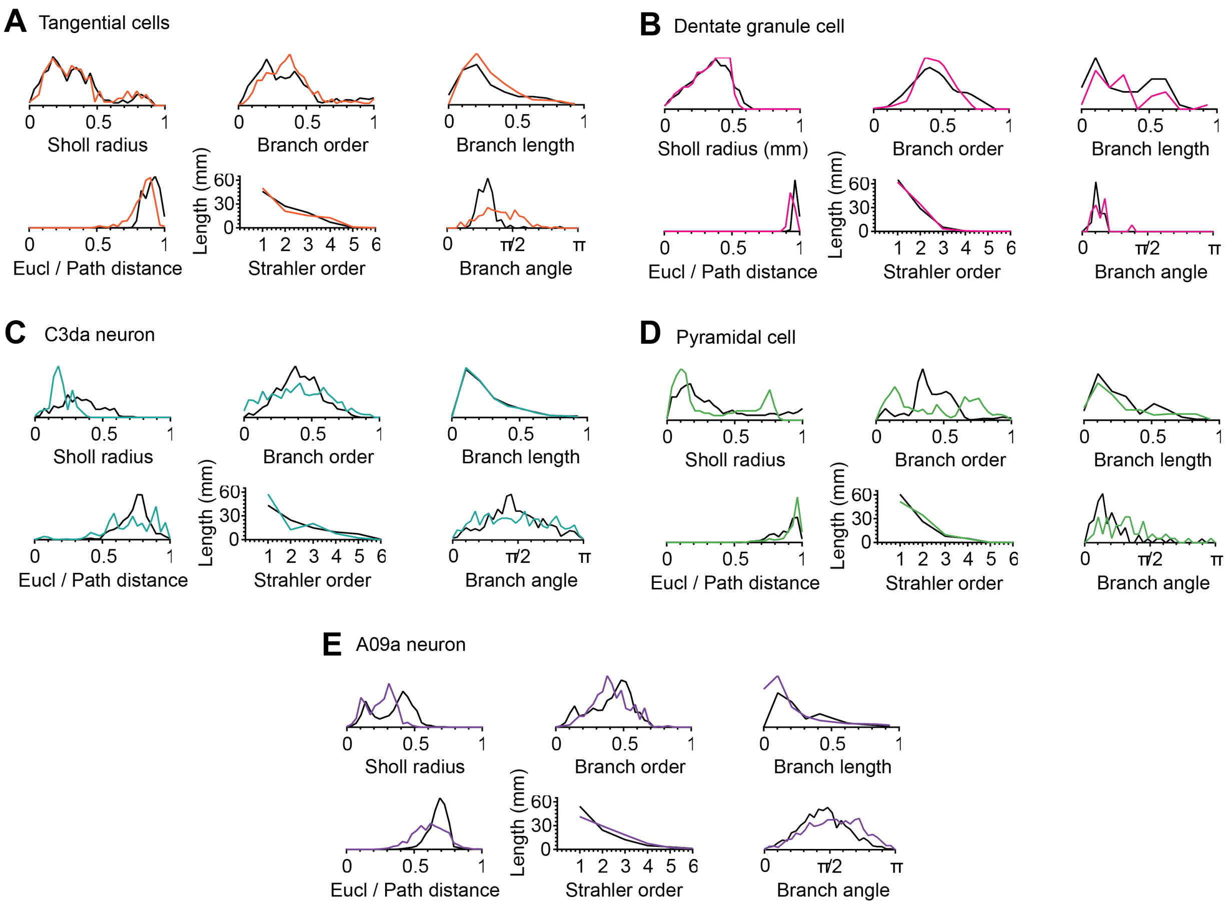


**Figure S7. Comparison statistics.**

Comparison statistics between the real (black lines) and synthesised models (coloured lines) in **Figure 6**: **A**, Tangential cells; **B**, Granule cells; **C**, C3da neuron **D**, Cortical pyramidal neuron; and **E**, L3 A90a basin neuron.

### Appendix

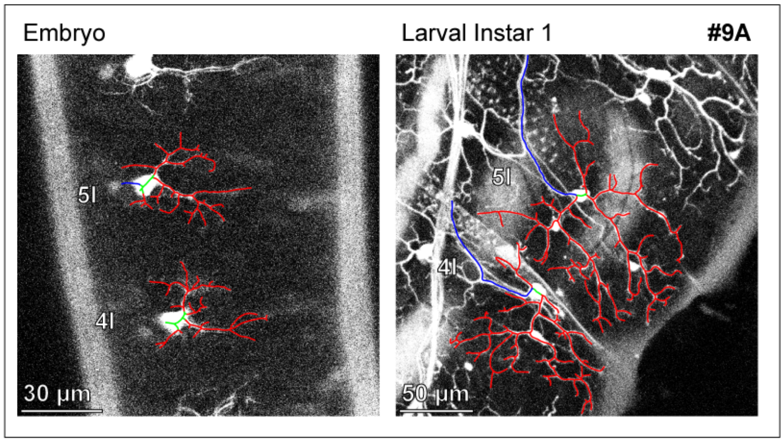
Embryo
Larval Instar 1
#9A
5l
4l
30 µm
5l
4l
50 µm

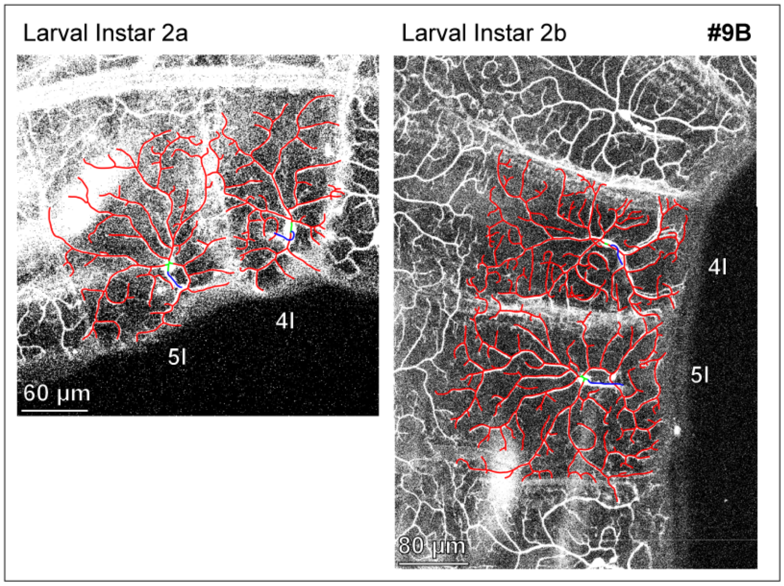
Larval Instar 2a
Larval Instar 2b
#9B
4l
5l
60 µm
4l
5l
80 µm

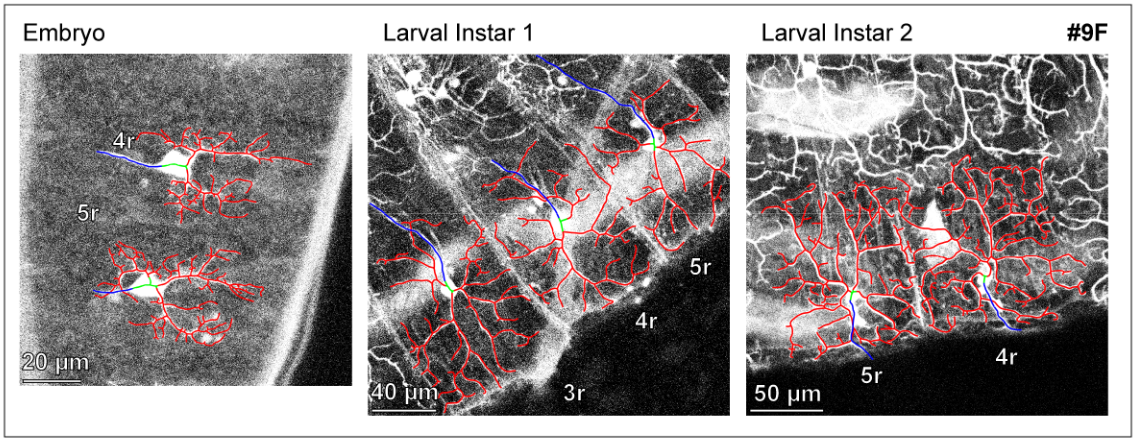
Embryo
Larval Instar 1
Larval Instar 2
#9F
4r
5r
20 µm
5r
4r
3r
40 µm
5r
4r
50 µm

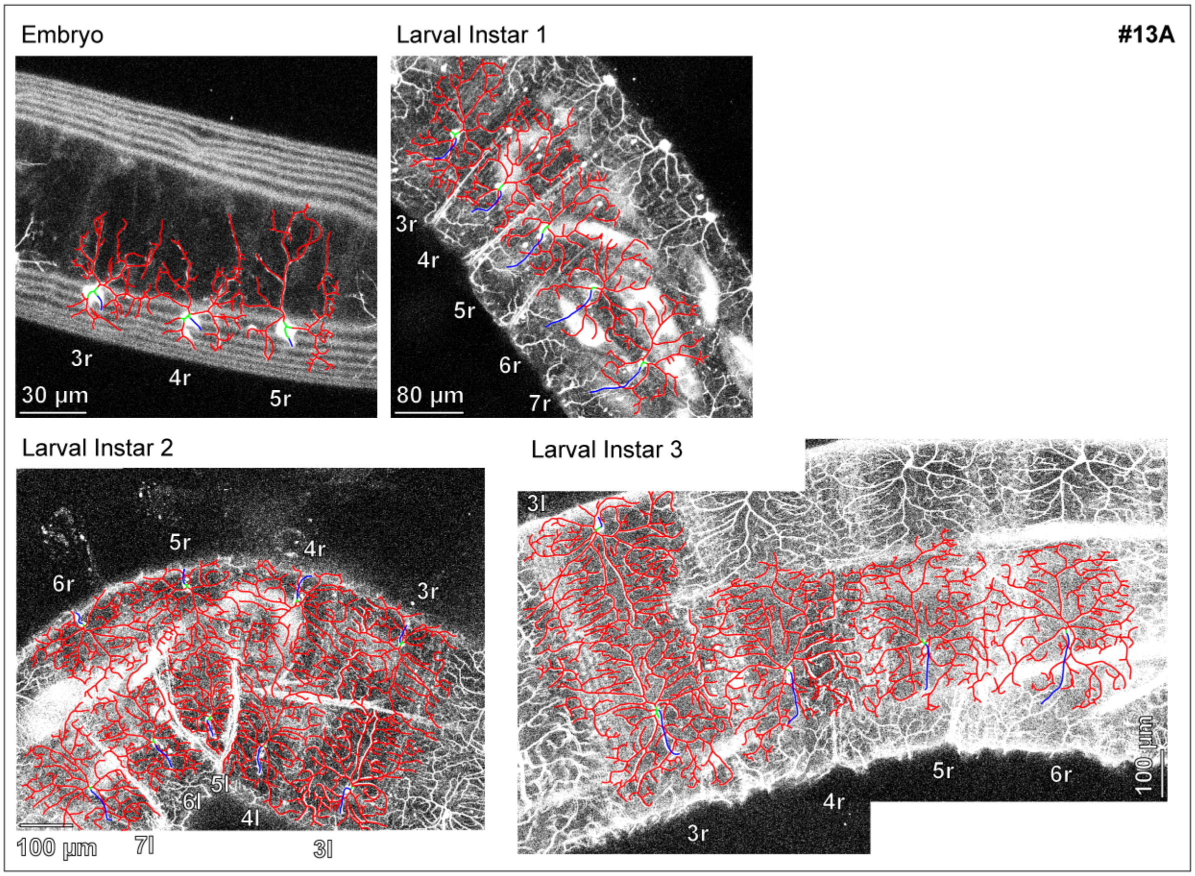
Embryo
Larval Instar 1
#13A
3r
4r
5r
30 μm
3r
4r
5r
6r
7r
80 μm
Larval Instar 2
5r
4r
6r
3r
5l
6l
4l
100 μm
7l
3l
Larval Instar 3
3l
5r
6r
4r
3r
100 μm

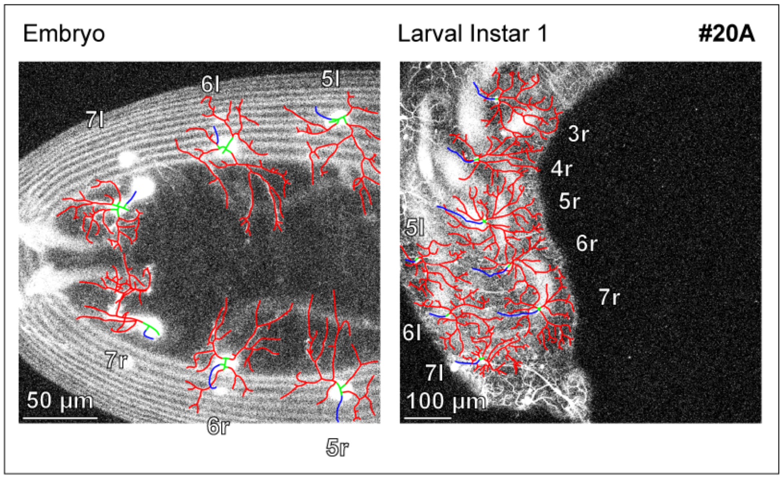
Embryo
Larval Instar 1
#20A
6l
5l
7l
7r
50 μm
6r
5r
3r
4r
5r
6r
7r
5l
6l
7l
100 μm

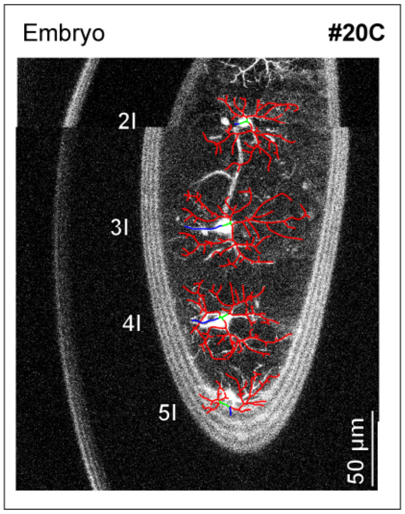
Embryo
#20C
2l
3l
4l
5l
50 μm

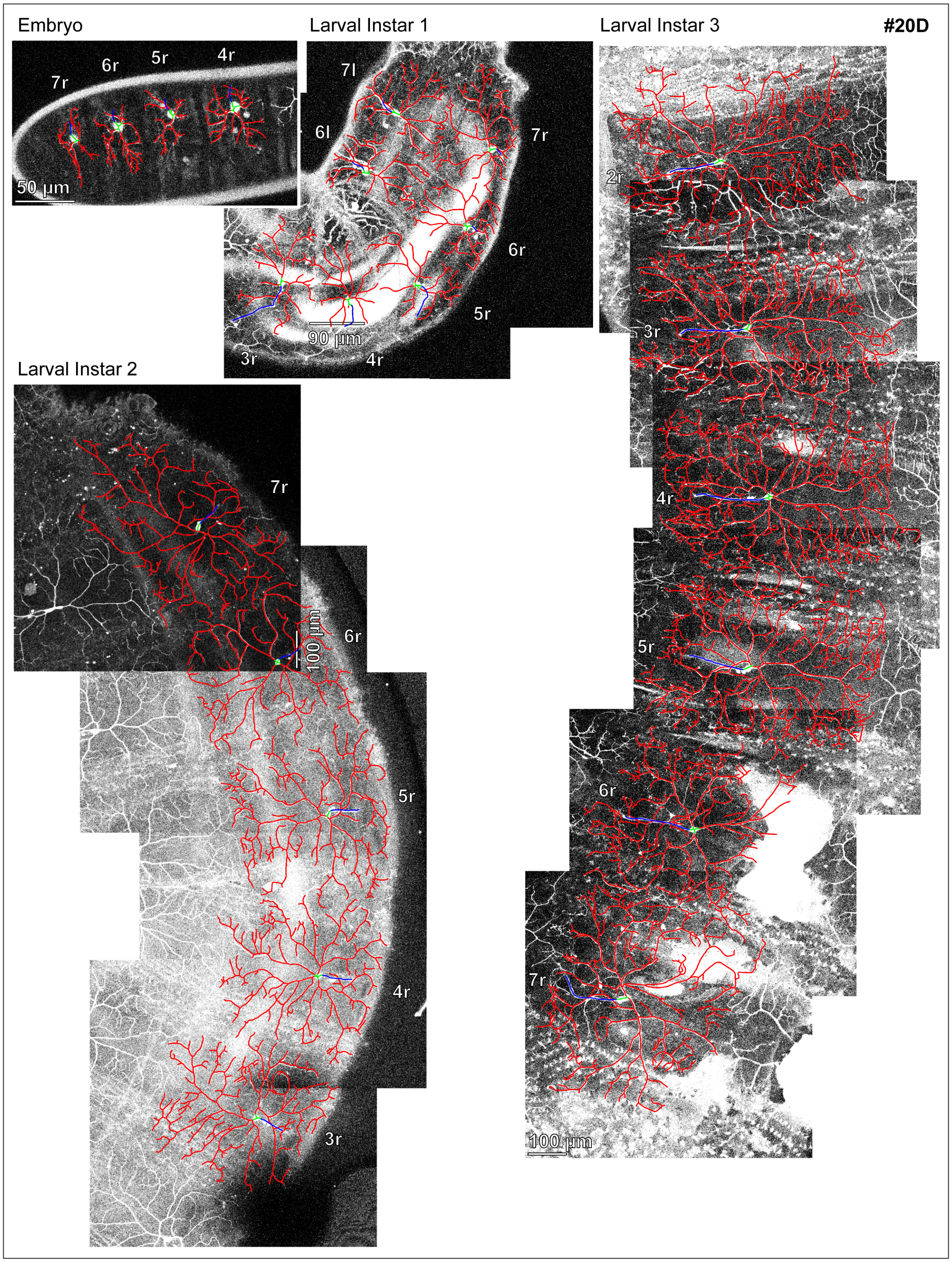

Embryo
Larval Instar 1
Larval Instar 3
#20D
7r
6r
5r
4r
50 µm
7l
6l
7r
6r
5r
90 µm
3r
4r
2r
3r
Larval Instar 2
7r
6r
100 µm
5r
4r
3r
4r
5r
6r
7r
100 µm

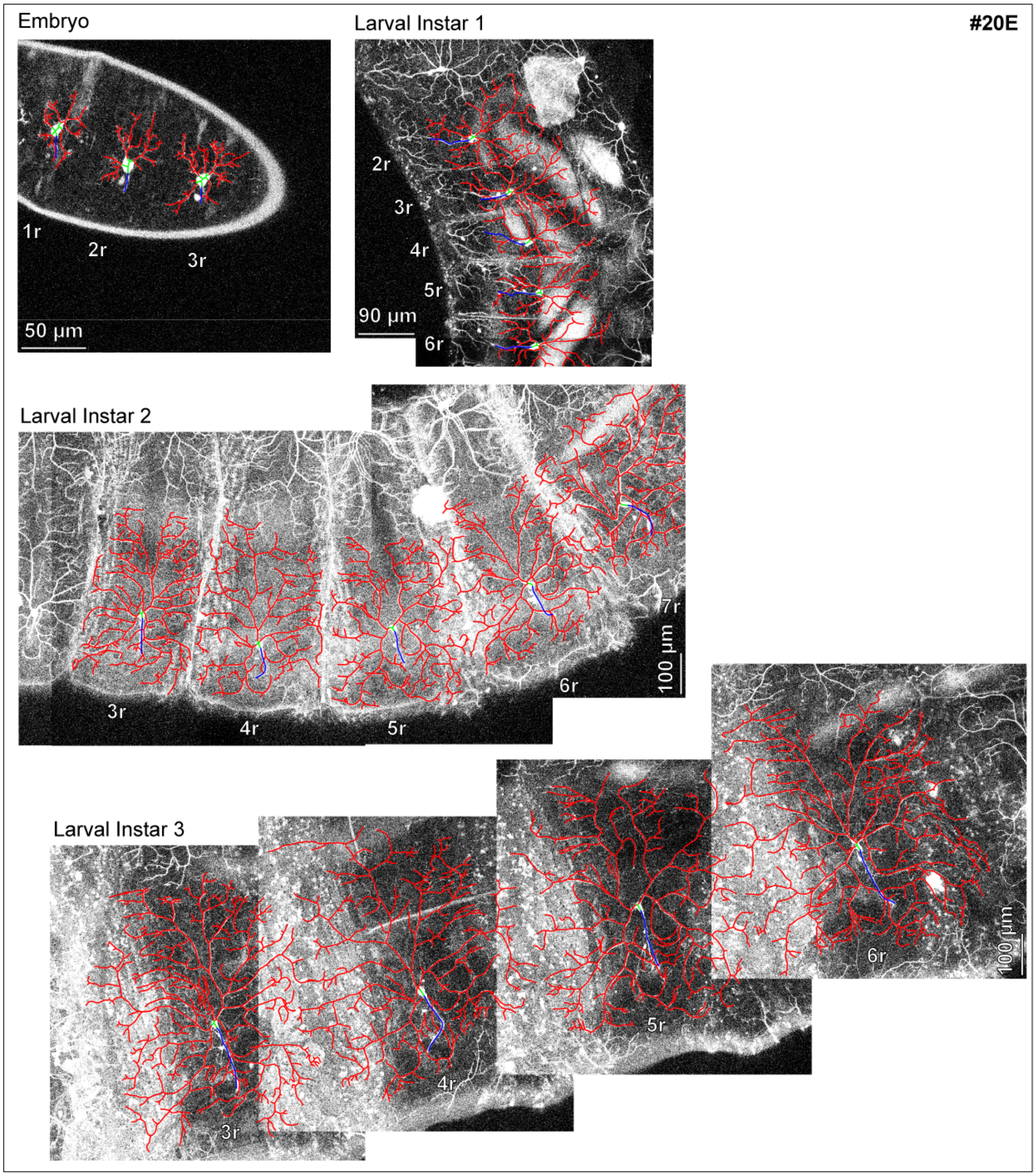
#20E
Embryo
1r
2r
3r
50 µm
Larval Instar 1
2r
3r
4r
5r
6r
90 µm
Larval Instar 2
3r
4r
5r
6r
7r
100 µm
Larval Instar 3
3r
4r
5r
6r
100 µm

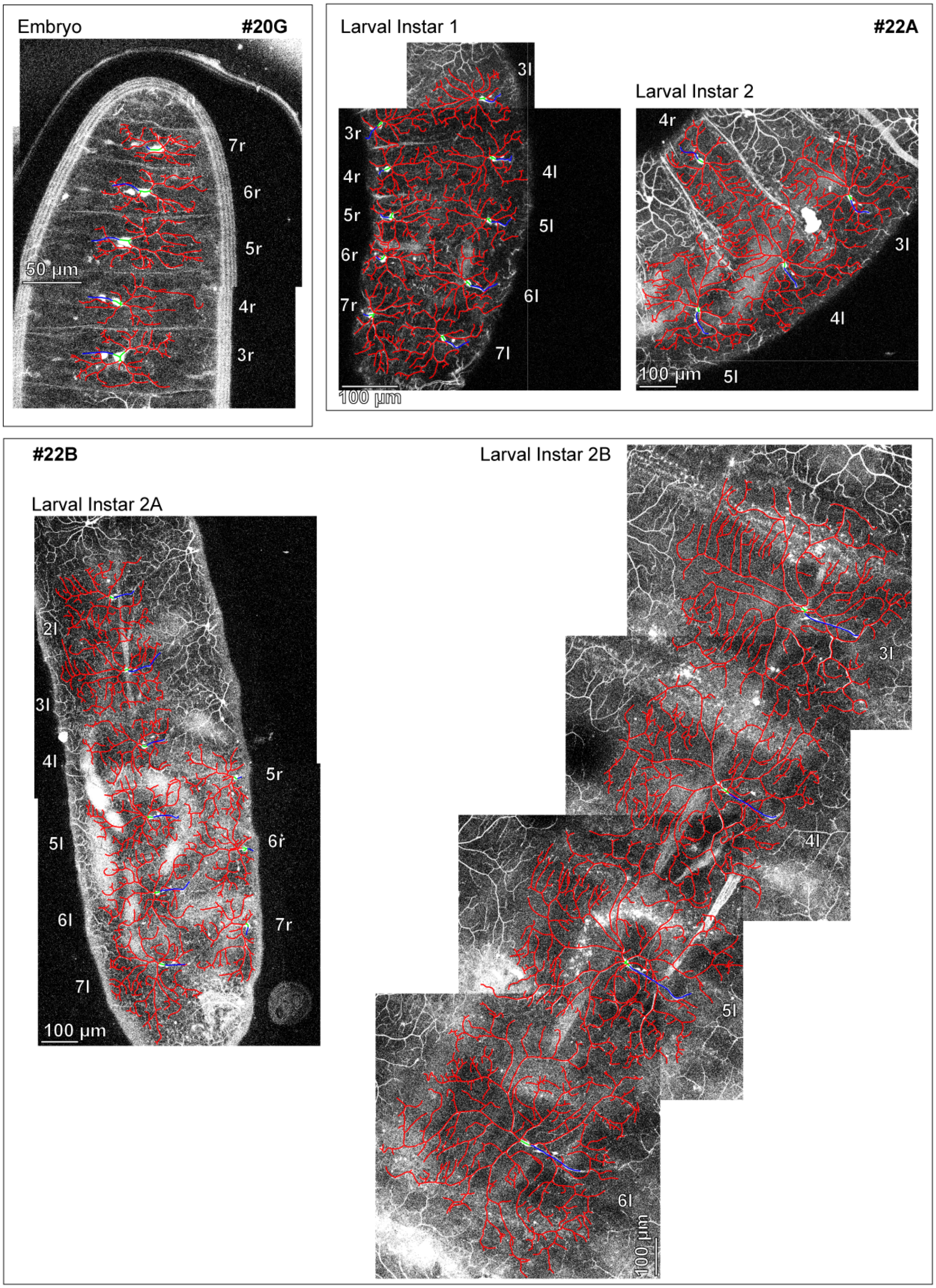
Embryo
#20G
7r
6r
5r
50 µm
4r
3r
Larval Instar 1
#22A
3l
3r
4r
4l
5r
5l
6r
6l
7r
7l
100 µm
Larval Instar 2
4r
3l
4l
100 µm
5l
#22B
Larval Instar 2B
Larval Instar 2A
2l
3l
4l
5r
5l
6r
6l
7r
7l
100 µm
3l
4l
5l
6l
100 µm

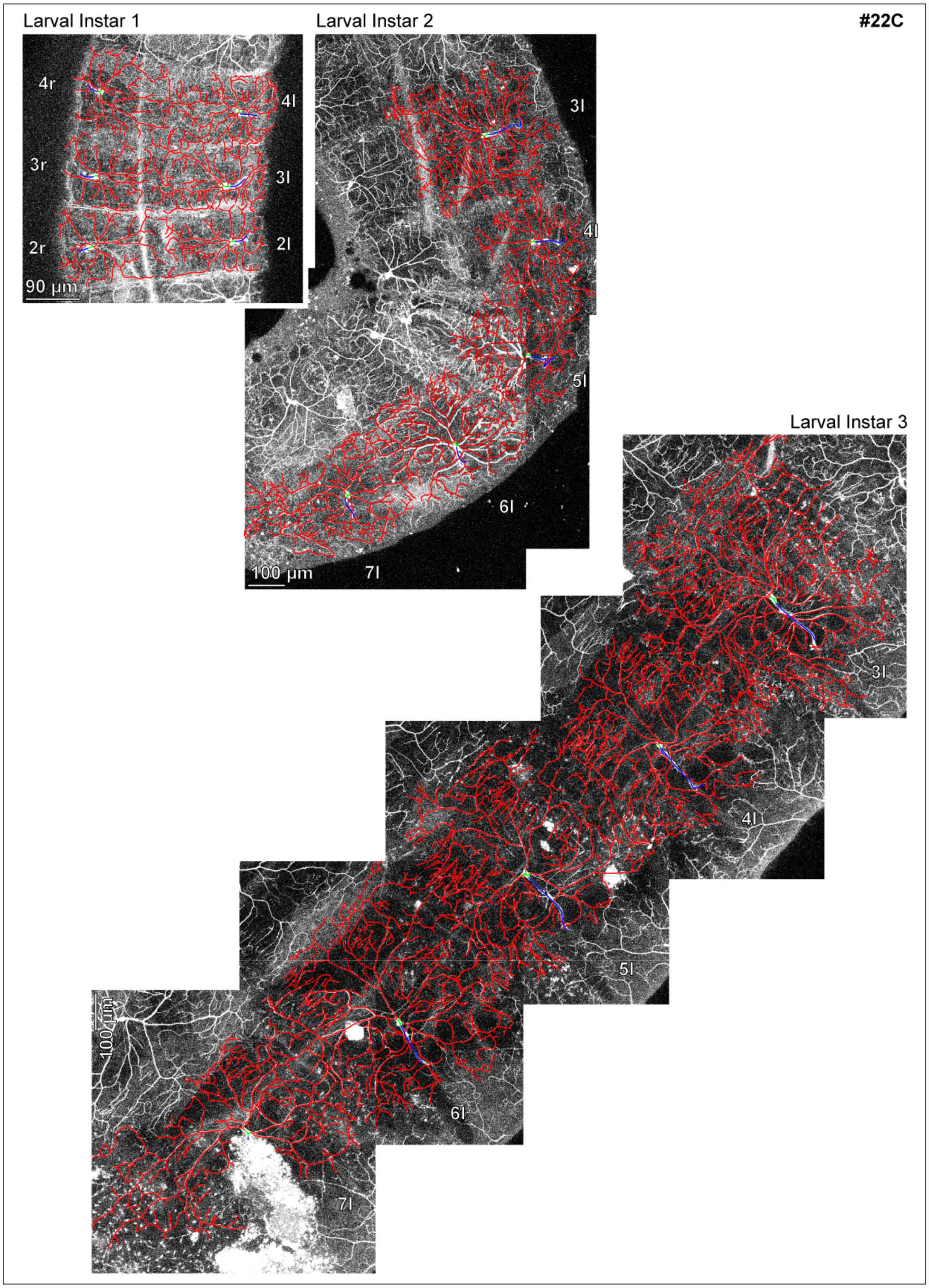
Larval Instar 1
Larval Instar 2
#22C
4r
4l
3r
3l
2r
2l
90 µm
3l
4l
5l
6l
7l
100 µm
Larval Instar 3
3l
4l
5l
6l
7l
100 µm

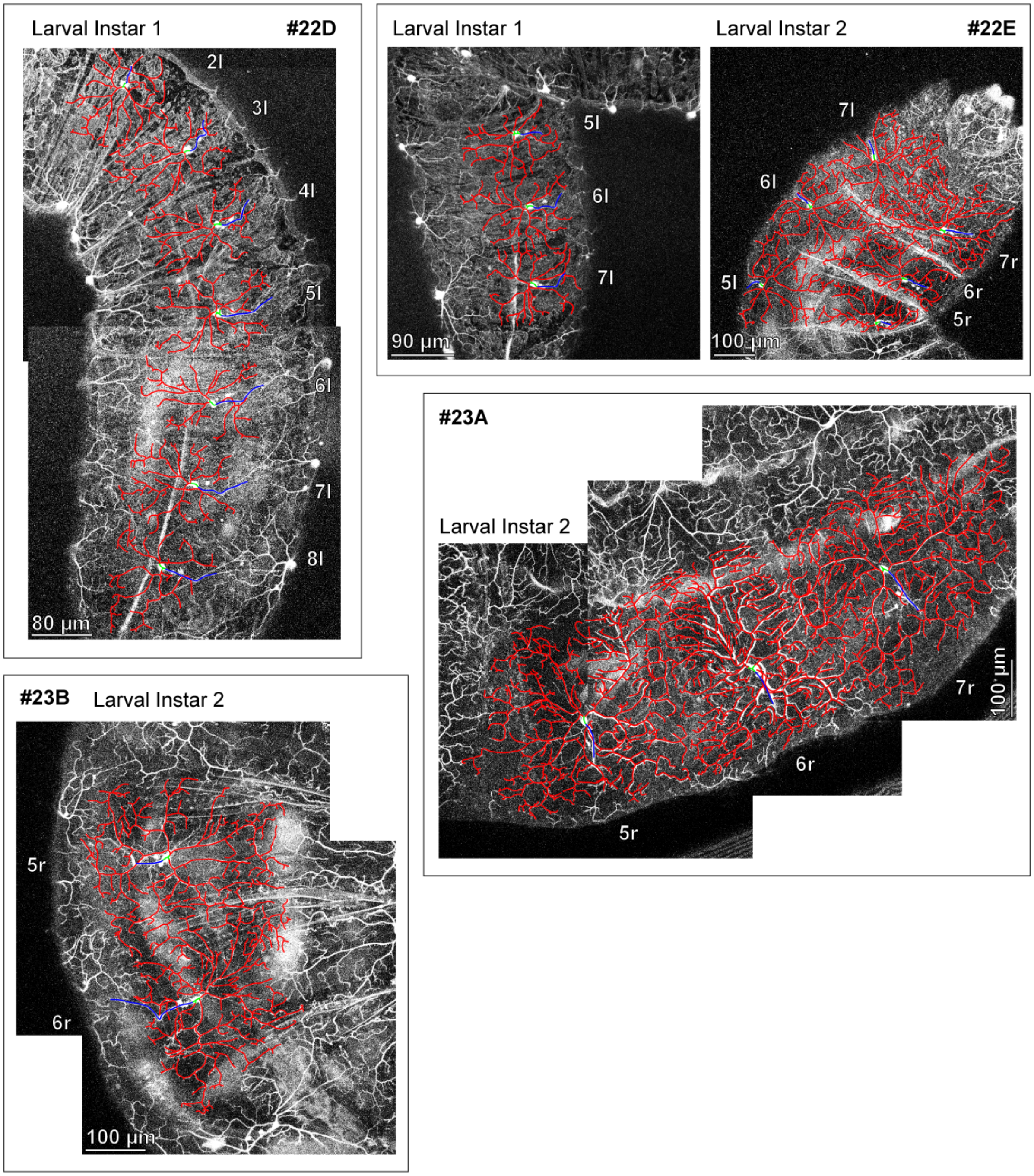
Larval Instar 1
#22D
2l
3l
4l
5l
6l
7l
8l
80 µm
Larval Instar 1
Larval Instar 2
#22E
5l
6l
7l
90 µm
7l
6l
5l
7r
6r
5r
100 µm
#23A
Larval Instar 2
7r
100 µm
6r
5r
#23B
Larval Instar 2
5r
6r
100 µm

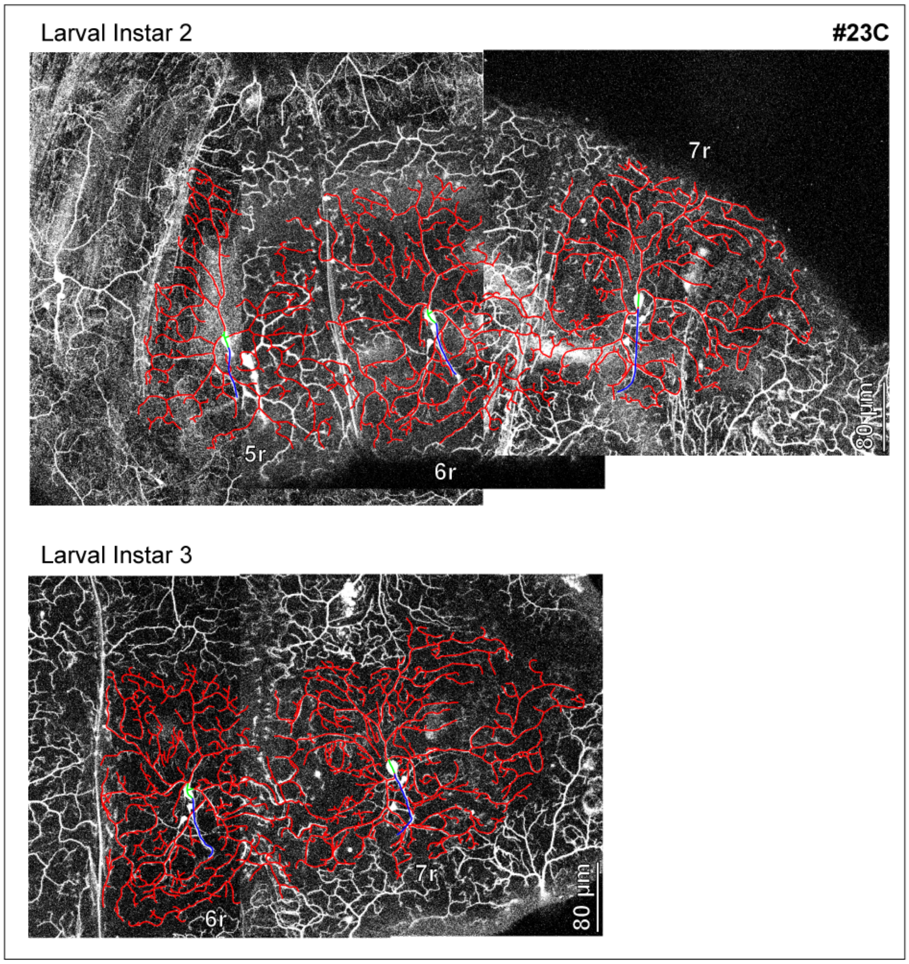
Larval Instar 2
#23C
7r
5r
6r
80 µm
Larval Instar 3
6r
7r
80 µm

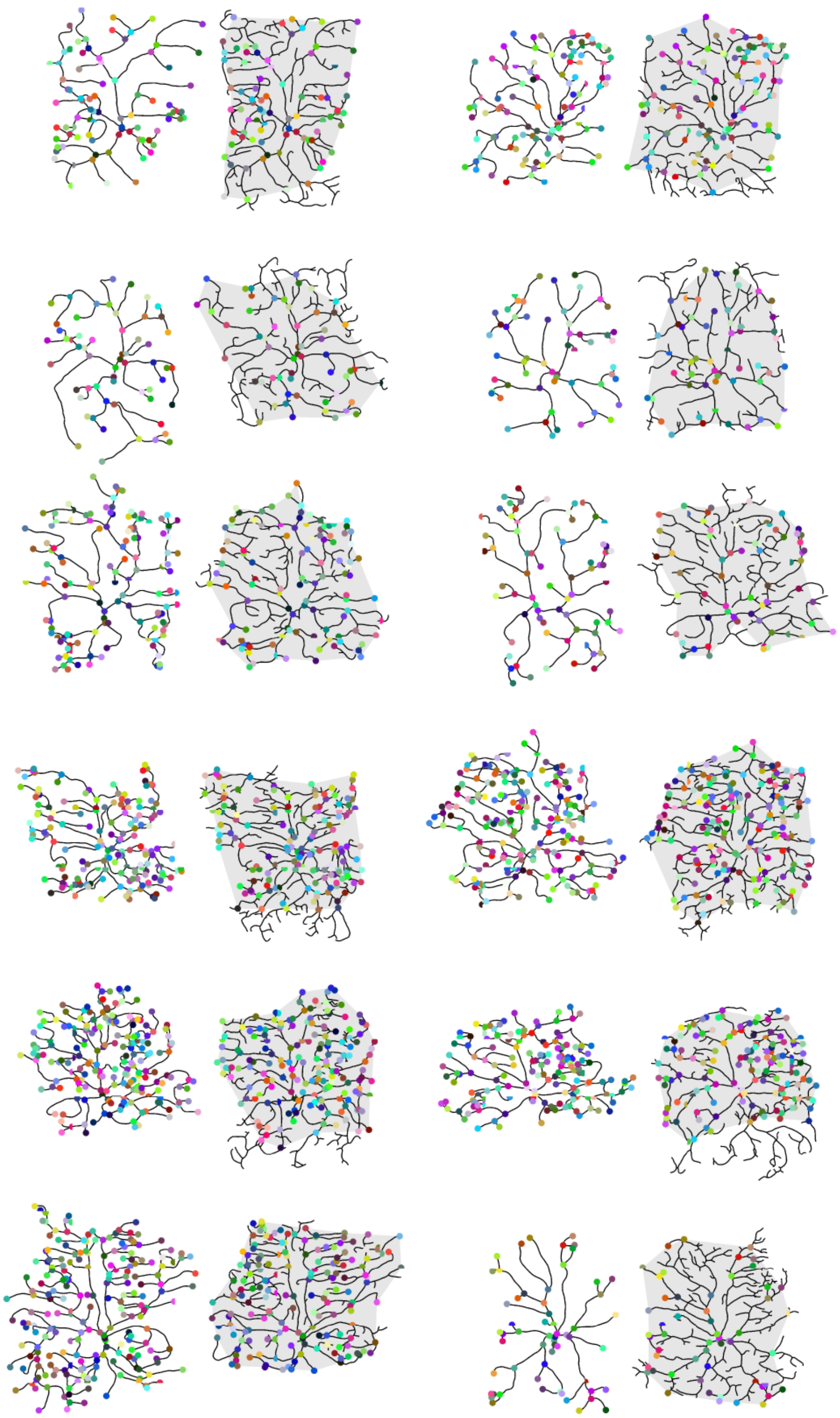

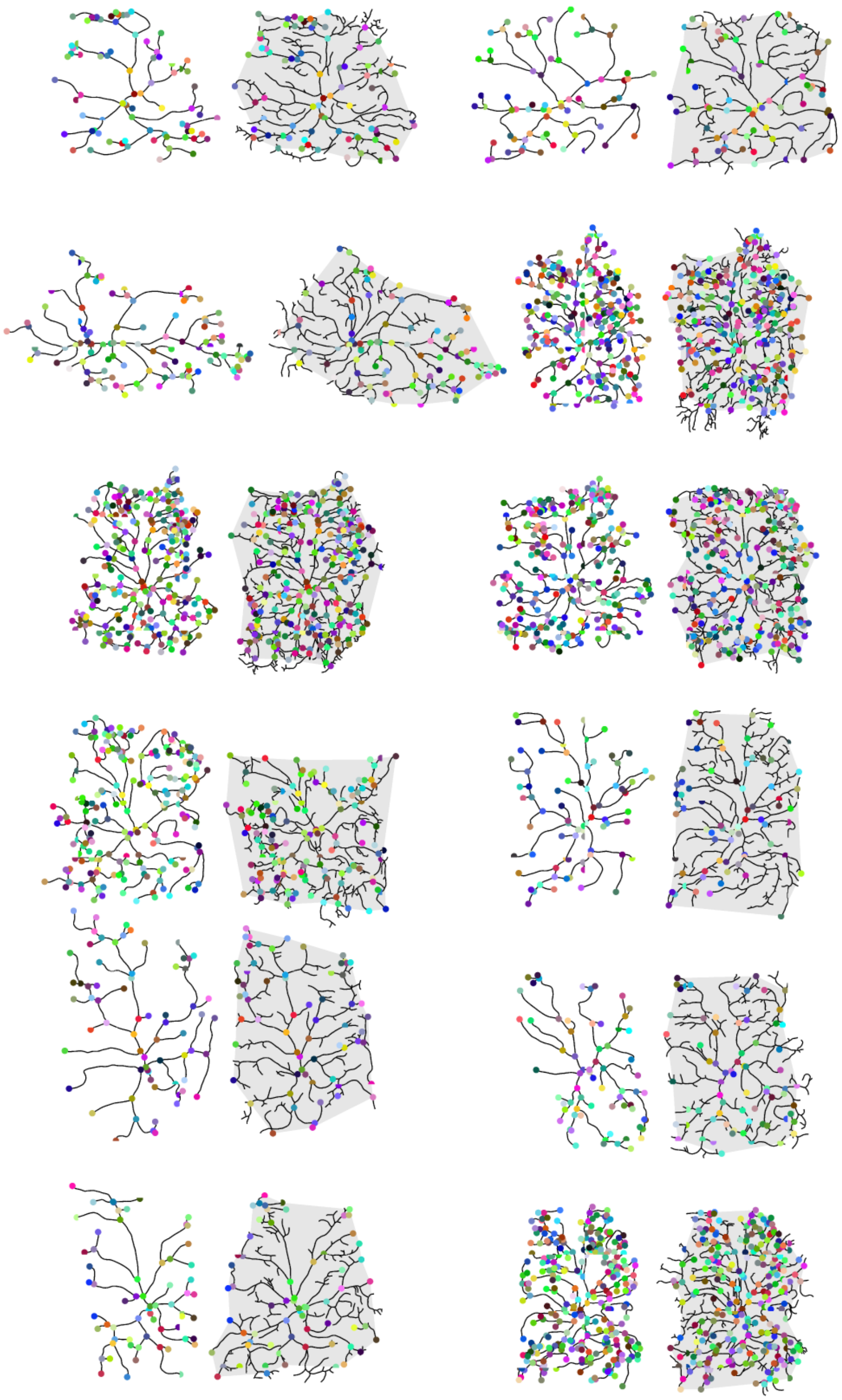

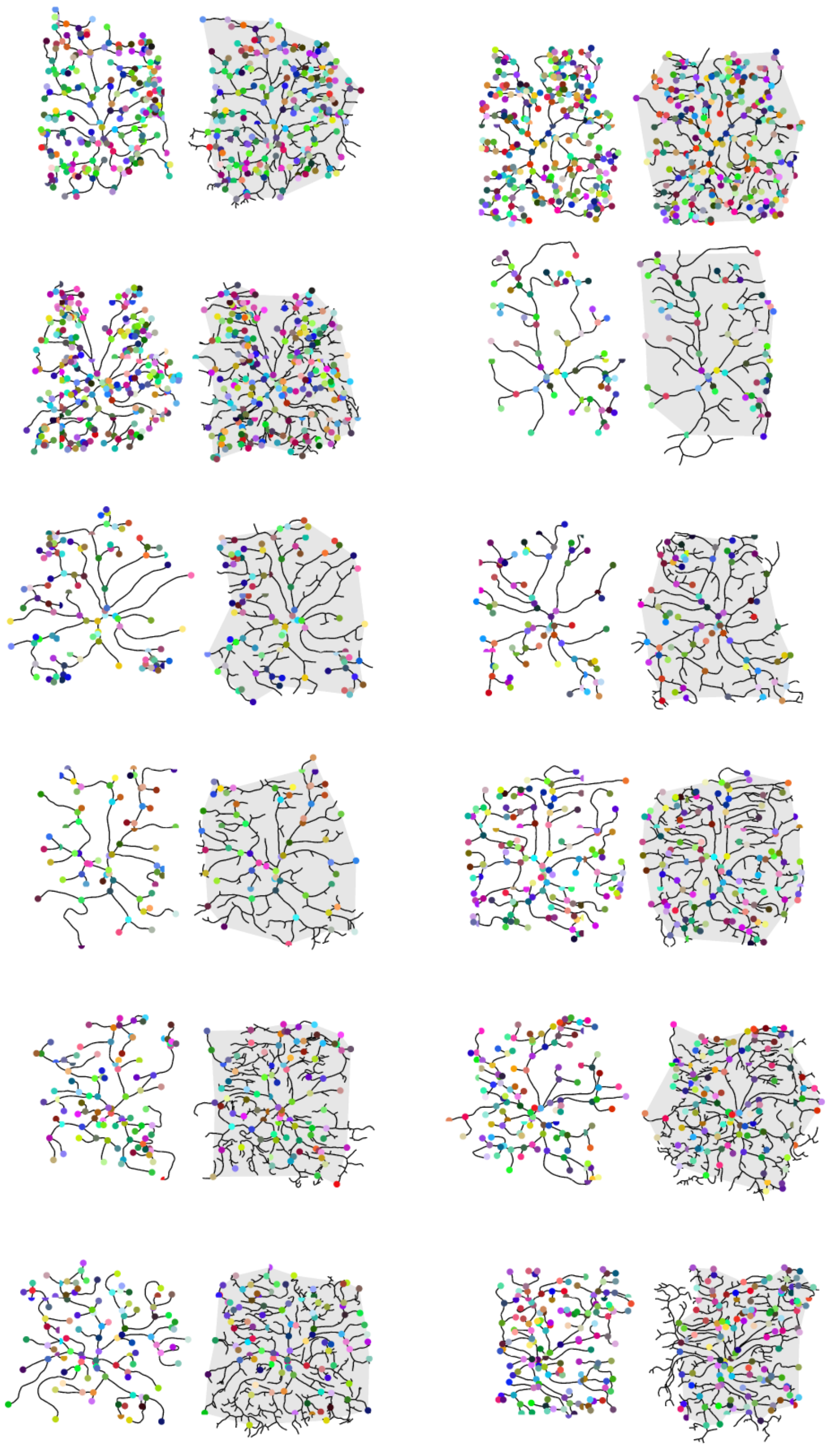

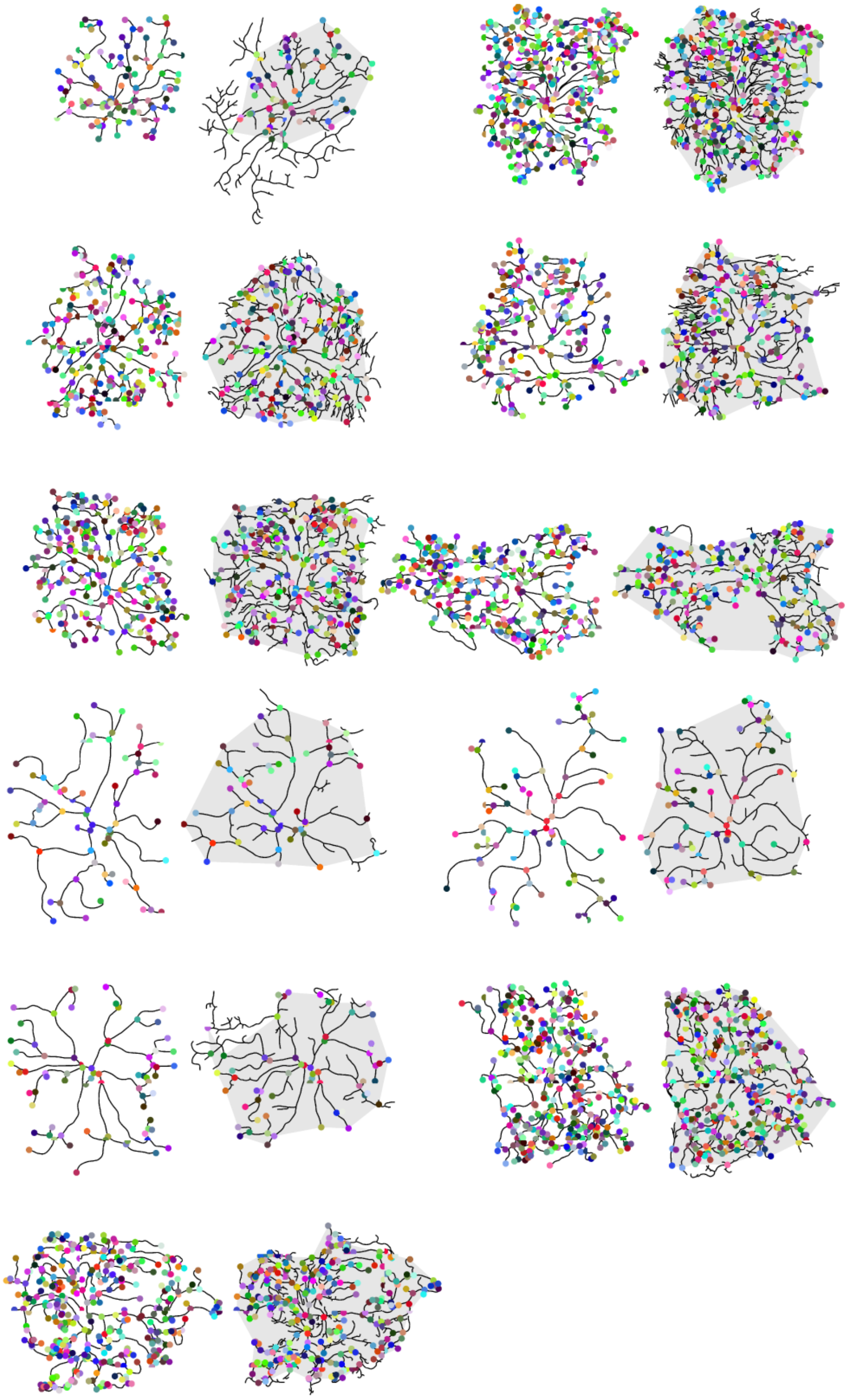
